# Tethering distinct molecular profiles of single cells by their lineage histories to investigate sources of cell state heterogeneity

**DOI:** 10.1101/2022.05.12.491602

**Authors:** Anna Minkina, Junyue Cao, Jay Shendure

## Abstract

Gene expression heterogeneity is ubiquitous within single cell datasets, even among cells of the same type. Heritable expression differences, defined here as those which persist over multiple cell divisions, are of particular interest, as they can underlie processes including cell differentiation during development as well as the clonal selection of drug-resistant cancer cells. However, heritable sources of variation are difficult to disentangle from non-heritable ones, such as cell cycle stage, asynchronous transcription, and measurement noise. Since heritable states should be shared by lineally related cells, we sought to leverage CRISPR-based lineage tracing, together with single cell molecular profiling, to discriminate between heritable and non-heritable variation in gene expression. We show that high efficiency capture of lineage profiles alongside single cell gene expression enables accurate lineage tree reconstruction and reveals an abundance of progressive, heritable gene expression changes. We find that a subset of these are likely mediated by structural genetic variation (copy number alterations, translocations), but that the stable attributes of others cannot be understood with expression data alone. Towards addressing this, we develop a method to capture cell lineage histories alongside single cell chromatin accessibility profiles, such that expression and chromatin accessibility of closely related cells can be linked via their lineage histories. We call this indirect “coassay” approach “THE LORAX” and leverage it to explore the genetic and epigenetic mechanisms underlying heritable gene expression changes. Using this approach, we show that we can discern between heritable gene expression differences mediated by large and small copy number changes, *trans* effects, and possible epigenetic variation.

## Introduction

Single cell molecular profiling technologies have revealed extensive gene expression heterogeneity, even between cells of a single cell type (Y. H. Choi & Kim, 2019; Li et al., 2022; Muto et al., 2021; O’Leary et al., 2020; Patel et al., 2014; SoRelle et al., 2021). Expression variation can arise from a number of sources, including transient phenomenon like cell cycle stage and transcriptional bursting (Tunnacliffe & Chubb, 2020), as well as stable genetic (Ben-David et al., 2018) or epigenetic (Bonasio et al., 2010) differences within a cell population. Stable sources of variation are of particular interest as they are “heritable” over multiple cell divisions, and can thus serve as substrates for selection, altering a cell population over time. Such heritable phenomena may underlie differentiation during normal organismal development as well as the acquisition of drug resistance in cancer (Salgia & Kulkarni, 2018). Yet within a set of single cell gene expression profiles, representing a population snapshot in time, it is difficult to distinguish between stable and transient expression variation. This is particularly challenging for cells of a single cell type, where transient differences may mask heritable variation when performing clustering analysis to distinguish cell states (Kiselev et al., 2019).

Heritable sources of expression variation have at least one property which distinguishes them from transient variation: because they are stable over multiple cell divisions, they should be shared by cells which are closely related by lineage. It follows that if all lineage relationships were known, we could discern heritable from non-heritable variation by assessing the distribution of variation across a lineage tree (**Figure 1a**). While transient variation should be randomly distributed, stably maintained expression states should cluster together within the tree, *i.e*. tracking to a common “founder” event. Thus, lineage histories, coupled to gene expression profiling, could potentially enable the differentiation of heritable vs. non-heritable sources of expression variation.

**Figure 1.**
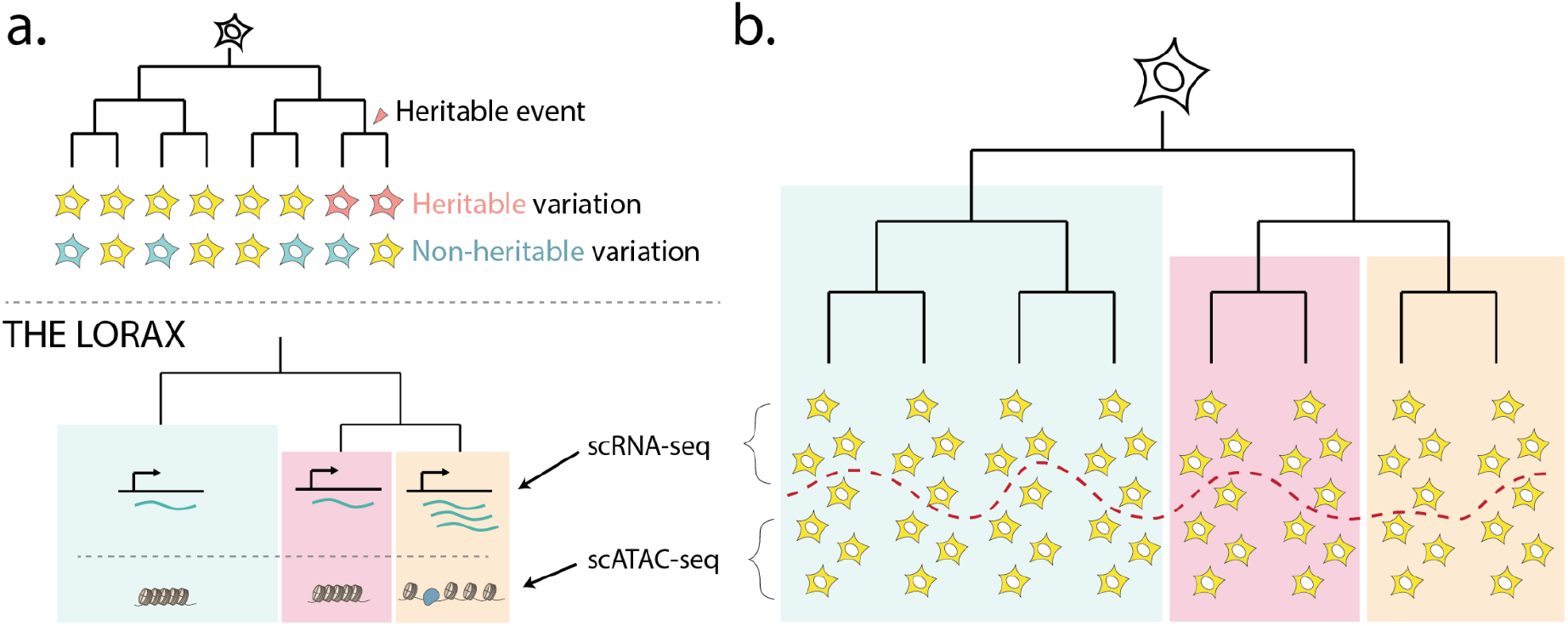
Tethering the molecular profiles of single cells by their lineage histories to investigate sources of cell state heterogeneity. (**a**) A framework to distinguish heritable from non-heritable sources of gene expression variation using lineage relationships. (**b**) A framework for tethering single cell expression (scRNA-seq) and chromatin accessibility (scATAC-seq) measurements via lineage relationships to investigate the mechanisms underlying heritable expression variation (THE LORAX).

Molecular methods for cell lineage history profiling compatible with concurrent expression profiling involve either static or progressive genetic barcoding. The static approach introduces short, transgenic barcodes to proliferating cells, such that closely related descendants share a barcode sequence (Biddy et al., 2018; Guo et al., 2019; Rodriguez-Fraticelli et al., 2018; Weinreb et al., 2020). Static barcoding might reveal heritable sources of gene expression that were acquired close to the time of labeling, but would presumably miss those occurring substantially earlier or later. In contrast, progressive lineage tracing methods (*e.g*. GESTALT and related methods), wherein cells accumulate sequence diversity at multiple genomic locations over time, facilitate reconstruction of multi-tier lineage trees, and might therefore be more sensitive with respect to detecting heritable gene expression variation (Alemany et al., 2018; Bowling et al., 2020; Chan et al., 2019; Hwang et al., 2019; Kalhor et al., 2017, 2018; Loveless et al., 2021; McKenna et al., 2016; Perli et al., 2016; Raj, Gagnon, et al., 2018; Raj, Wagner, et al., 2018; Spanjaard et al., 2018; Wagner et al., 2018).

A high diversity of labels can be achieved via CRISPR/Cas9, where imperfect double strand break repair via NHEJ can generate a variety of outcomes (referred to here as “edits” or “indels”) (Alemany et al., 2018; Bowling et al., 2020; Chan et al., 2019; Kalhor et al., 2017, 2018; Loveless et al., 2021; McKenna et al., 2016; Perli et al., 2016; Raj, Gagnon, et al., 2018; Raj, Wagner, et al., 2018; Spanjaard et al., 2018; Wagner et al., 2018). Over many cell divisions, the pattern of indels that accumulate at CRISPR/Cas9 targets are informative with respect to the lineage relationships amongst the cells in which they occur. Most strategies reported to date, whether implemented *in vitro* or *in vivo*, place several targets in tandem, such that the edits at these multiple targets can be recovered within a single DNA or RNA-derived sequencing read (Alemany et al., 2018; Bowling et al., 2020; Chan et al., 2019; Kalhor et al., 2017, 2018; Loveless et al., 2021; McKenna et al., 2016; Perli et al., 2016; Raj, Gagnon, et al., 2018; Raj, Wagner, et al., 2018; Spanjaard et al., 2018; Wagner et al., 2018).

In practice, however, there are a number of technical issues that limit this approach. First, arrays of CRISPR/Cas9 targets frequently acquire large deletions when concurrent DSBs at different targets within the array are joined, potentially excising previously recorded information at intervening targets. Second, read length limitations require targets to be placed close to one another, such that the editing of one target risks corrupting adjacent targets. Third, although it is possible to capture CRISPR/Cas9-edited lineage targets as part of a single cell RNA-seq (scRNA-seq) profile, this has usually been inefficient in practice. For example, using InDrops to capture a tandem array of 10 CRISPR targets alongside single cell transcriptomes in juvenile zebrafish brains, Raj *et al*. (2018) recovered lineage profiles from just 6-28% of cells with expression profiles (Raj, Wagner, et al., 2018). Similarly, using 10X Genomics to capture arrays of 3 CRISPR targets from mouse embryos alongside scRNA-seq (3-15 array integrations per embryo), Chan *et al*. (2019) recovered at least one edited lineage array from 15-75% of cells per embryo, but just one target array was captured efficiently (>25% of cells) in 6 of 7 embryos (Chan et al., 2019). In each case, both target design and the method of capturing lineage targets during scRNA-seq likely contributed to the limited recovery.

Here, we introduce a CRISPR-based lineage tracing approach in which many distinct lineage recording loci are integrated independently throughout the genome. These targets can each accommodate relatively large deletions and insertions. We further show that, with targeted enrichment, they can be captured efficiently alongside transcriptomes via a combinatorial indexing approach (sci-RNA-seq) (Cao et al., 2017, 2019). To analyze data generated from a proof-of-concept *in vitro* monoclonal expansion, we developed a lineage tree reconstruction algorithm that is robust to missing data and recurrences (*i.e*. where identical edits occur independently), and validate the algorithm using copy number alterations (CNAs) that are evident in expression data. We show that incorporating lineage relationships into expression analysis reveals abundant heritable expression variation, including instances that are clearly explained by CNAs, but also many which are not.

Finally, towards investigating the mechanism(s) underlying expression heritability, we develop an approach to capture cell lineage relationships alongside single cell chromatin accessibility. We show that we can link two distinct molecular features—gene expression and chromatin accessibility—via their lineage profiles (**Figure 1b**). We then use these lineage-tethered features to further distinguish between expression changes which can be explained directly by copy number alterations, ones likely mediated by *trans* effects of copy number alterations, and ones which are more likely to have resulted from a stable change in *cis* regulatory state. We term this approach THE LORAX: Tracking Heritable Events via Lineage-based Ordering of chRomatin Accessibility & eXpression profiles.

## Results

### Concurrent profiling of many independent CRISPR lineage targets and gene expression via single cell combinatorial indexing

We first set out to design a CRISPR/Cas9-based lineage tracing strategy that addresses outstanding technical challenges. Reconstructing an accurate, multi-tier lineage tree from progressively acquired edits requires the following: (a) multiple editable loci such that successive tagging can occur in a single lineage over time; (b) a high probability of diverse editing outcomes at a single target, such that identical edits at that target are unlikely to occur independently in different cells; (c) controllable editing machinery, such that target capacity is not exhausted quickly after editing onset; (d) permanence of edits, such that they are not likely to be overwritten or lost; and (e) a high rate of capture of editing information alongside single cell profiling of other features. Towards realizing these features, we designed a construct in which individual targets are integrated independently across the genome and captured as separate transcripts (**Figure 2a-b**). Each target contains a unique identifier sequence, which is positioned such that the target can accommodate up to a 70 bp deletion centered at the cut site without corrupting the identifier, as well as, assuming 300 bp read lengths, insertions of up to 105 bp. The sgRNAs are delivered on the same lentiviral construct as the targets, with targets expressed from a highly active EF-1α promoter to enable lineage capture from mRNA.

**Figure 2.**
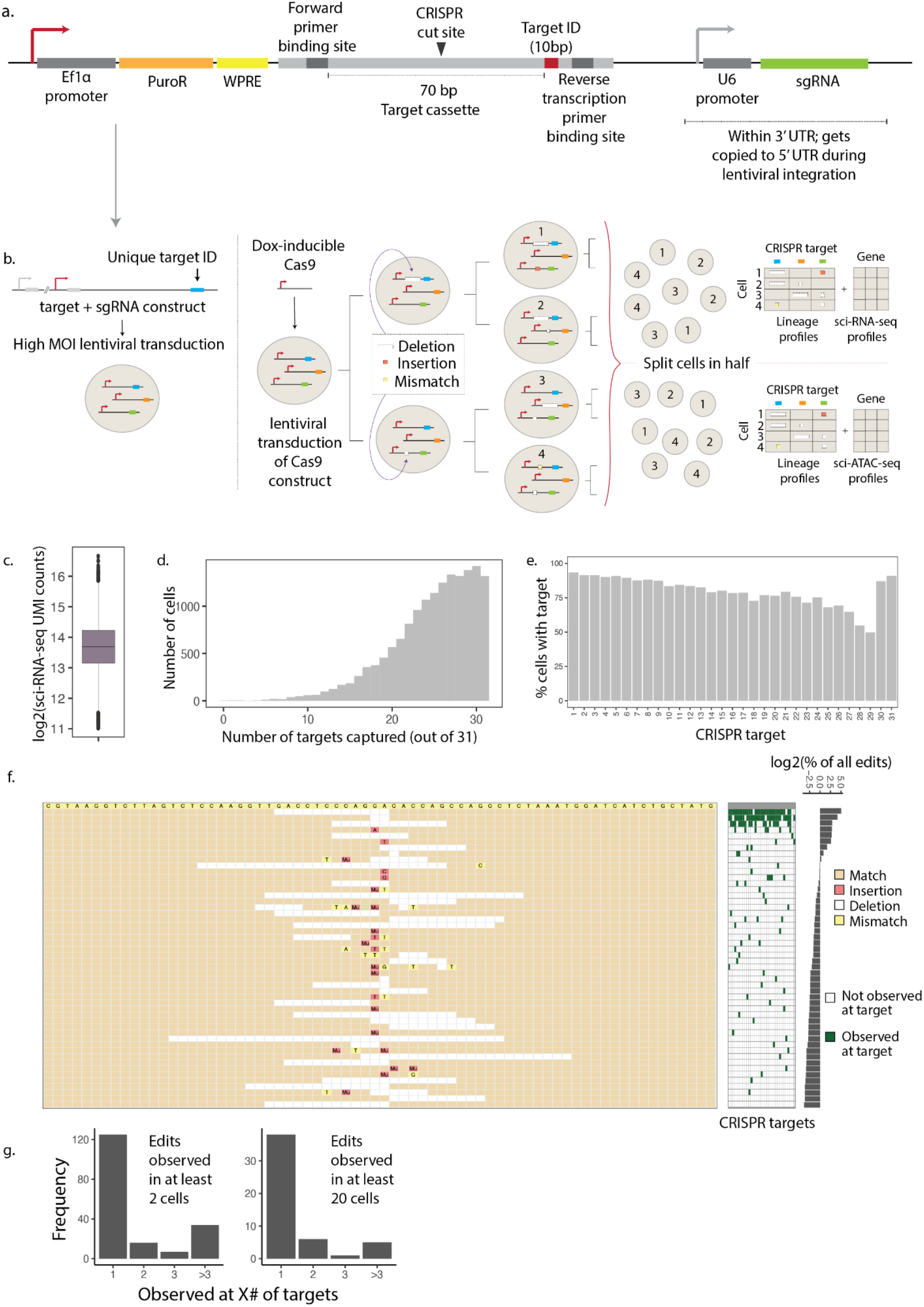
Experimental design, target capture rate and CRISPR editing diversity. (**a**) Target vector design. A target cassette was integrated into the CROP-seq vector (Datlinger et al., 2017) as shown. (**b**) Schematic of experimental workflow. Cells were transduced at high MOI with constructs containing an sgRNA and barcoded target sequences, such that many integration events per cell were expected. A single clone was then transduced with a doxycycline-inducible Cas9 vector, single cells were sorted, and a single founder cell was allowed to divide for 35 days while editing occurred. The final cell population was split for either target capture alongside sci-RNA-seq or sci-ATAC-seq. (**c**) Log-scaled boxplot of UMI counts for sci-RNA-seq (not including enriched target UMIs). Box shows median and encompasses counts in the second and third quartiles. Whiskers depict the interquartile range, with outliers shown. (**d**) Histogram of the number of targets captured per cell. (**e**) Percent of cells from which each individual target was captured. Targets 30 & 31 were duplicated (see text), and hence artificially appear to have a high rate of capture. (**f**) Left: Top 50 most abundant editing patterns. Insertions are shown one base left of the insertion site; “Mu”: multi-base insertion. Middle: Targets at which the editing pattern is observed in at least 20 cells. Right: Log-scaled percentage of all edits represented by the top 50 editing patterns. (**g**) Proportion of editing patterns observed in 1, 2, 3, or more than 3 targets, if considering editing patterns appearing in at least 2 cells at a single target (left), or at least 20 cells (right).

To generate cells with a high capacity for lineage recording, we transduced HEK293 cells at a high multiplicity-of-infection (MOI) with this construct and attempted to establish clonal populations. Even in the absence of editing, most clones grew poorly, with the lentiviral integrations themselves at this high MOI potentially contributing to toxicity. Across 26 clones, we observed integration counts ranging from 2 to 53, with a median of 11 integrations (**Supplementary Fig. 1a**). We moved forward with a robust clone bearing 36 unique integrations, as evidenced by the diversity of unique identifier sequences (“target IDs”; **Supplementary Fig. 1b**). To induce editing, we transduced this clone again with a doxycycline-inducible Cas9 lentiviral construct, sorted single cells, and allowed a clonal population to grow from a single founder cell (such that all progeny cells comprise a single lineage tree). Interestingly, only 32 unique target IDs were observed after this second round of cloning, potentially due to karyotypic instability (discussed further below), while one integrant contained a mutation that corrupted its target site (**Supplementary Fig. 1b**).

After 35 days of expansion of this clone, with passaging as needed (**Methods**), a portion of the cells were harvested for single cell expression and lineage analysis, while the remaining cells were frozen down for subsequent profiling of chromatin accessibility. Of note, although doxycycline was not applied, we nonetheless observed diverse and progressive editing with this clone, presumably because of leaky expression of Cas9 (Costello et al., 2019). For concurrent acquisition of whole cell transcriptomes alongside lineage information, we performed 96 × 768 sci-RNA-seq, with processing of cells in eight batches during the second indexing step (Cao et al., 2017, 2019). To facilitate the efficient recovery of lineage targets from each cell, we introduced a supplemental set of reverse transcription primers during the first round of indexing, and split the material in half prior to indexed PCR during the second round of sci-RNA-seq2, with one half being used for the general transcriptome, and the other half for targeted recovery of the lineage profiles (**Methods**).

These libraries were sequenced, and the resulting reads were adaptor-trimmed, aligned to the reference human genome, and deduplicated. For the single cell transcriptomes, we observed a median of 13,212 UMIs per cell, across 15,525 cells (**Figure 2c**). For the 31 retained, uncorrupted lineage targets (**Supplementary Fig. 1b**) each bearing a unique target ID sequence in the resulting reads, we observed a high rate of capture, with ≥ 25 captured from 59% of cells, ≥ 20 from 85% of cells, and ≥ 10 from 99% (**Figure 2d**). Target capture rates were unevenly distributed across the eight batches of indexed PCR amplification, likely due to slight technical differences (**Methods**; **Supplementary Figure 2a-b**). Recovery varied across the integrations as well, with each target ID recovered in a median of 80% of cells (range 50% to 93%) (**Figure 2e**), presumably due to position effect variegation and/or early karyotypic instability or large deletions associated with more frequently lost targets. Overall, these results indicate that a modified version of sci-RNA-seq can be used to efficiently recover transcriptomes alongside dozens of lineage target integrants from each of many single cells.

We next performed a series of filtration steps, removing cells with limited lineage information as well as those deemed likely to be doublets. First, cells were filtered to those with at least 10 lineage targets recovered, at least one of which was edited. In some cases, an edit could not be resolved, as more than one editing pattern seemed to exist for a given lineage target integrant (**Methods**). We termed these edits “ambiguous.” Cells associated with more ambiguous than unambiguous edits, presumably doublets, were removed, as were cells with excessively high UMI counts (**Methods**; **Supplementary Fig 2c-d**). The single cell transcriptomes and associated lineage targets of the remaining 10,234 cells were carried forward for all subsequent analyses.

Across this entire dataset, we observed 461 unique editing patterns of the common target sequence, of which 182 were independently observed in at least 2 cells in association with the same target ID. The remainder may correspond to real events that occurred late in the expansion and were thus only sampled once, or alternatively PCR or sequencing errors. The 50 most frequently observed edits, across all cells and target IDs, are shown in **Figure 2f**. Of note, edits that recur independently as well as edits that occurred early during clonal expansion will both appear “common” by this measure. The three most frequently observed edits, together comprising 58% of all edits, appear to be recurrent: they occur in association with the majority of target IDs (**Figure 2f**), and furthermore correspond to outcomes anticipated to be favored by microhomology (Sfeir & Symington, 2015). Such frequent editing outcomes complicate tree construction, and can be avoided in the future through better target design (W. Chen et al., 2019). However, the clear majority of editing outcomes were only observed in association with a single target ID, consistent with their origination from a single event during the clonal expansion (**Figure 2g**).

Unexpectedly, two targets (#30 & #31) contained a large number of ambiguous editing calls—two distinct editing patterns convincingly present in association with the same target ID in the same single cell. This is consistent with a duplication event, *i.e*. in which the locus in which the target ID resides was duplicated early in the clonal expansion, or more likely during the second round of cloning. Additional evidence, discussed further below, of large-scale CNAs in the transcriptome data, corroborates this hypothesis. Rather than filtering out these targets, we “duplicated” them *in silico*, parsimoniously distributing the top two edits associated with these target IDs in a given cell, while minimizing the number of independent editing events required to explain them (**Methods**). As such, in the end, single cell lineage profiles contained 33 unique targets.

### Reconstructing lineage relationships using single cell lineage profiles

The reconstruction of cell lineage trees from CRISPR-edited targets has proven to be a difficult problem (Gong et al., 2021; Salvador-Martínez et al., 2019). Although phylogenetic reconstruction methods can in principle be applied here, several factors make this practically challenging. First, the amount of information within a lineage profile is limited to the number of targets that are edited and successfully recovered; the inefficient recovery observed in most studies to date results in substantial “missing data”. Second, recurrent events, *i.e*. the same edit occuring more than once independently at the same target, can be much more likely than in more conventional phylogenetic datasets, further complicating reconstruction. Third, it is computationally impractical to apply many popular phylogenetic algorithms to the large number of cells profiled with CRISPR-based lineage tracing, particularly those relying on generating a subset of all possible trees and choosing the most likely among them. To overcome this, one group employed a greedy approach to split cells into subgroups, generating subtrees of subgroups and merging them at the end (Jones et al., 2020). However, this approach was hindered by missing data in individual cell lineage profiles, which frequently split closely related cells across multiple subgroups.

On the other hand, CRISPR-based lineage tracing data has one feature which makes it more amenable to step-wise (rather than probabilistic) reconstruction strategies—the starting state of each target, *i.e*. unedited, is known. Given this, it is at least theoretically possible to employ a divisive, greedy approach to build a highly accurate tree (**Figure 3c,d**). In the proposed algorithm, all cells begin as a single group, which is split into two groups based on the presence vs. absence of the most common editing pattern associated with a single target. This edit is inferred by its frequency to have occurred earlier than other edits in cells belonging to the group. This splitting step is iterated on each sub-group, and each sub-sub-group, etc., terminating when all unique lineage profiles are represented by individual branches. Subsequently, unsupported bifurcations (those wherein a branch is not defined by a specific editing event(s)) are collapsed, such that more than two branches can arise from a single inferred ancestor.

**Figure 3.**
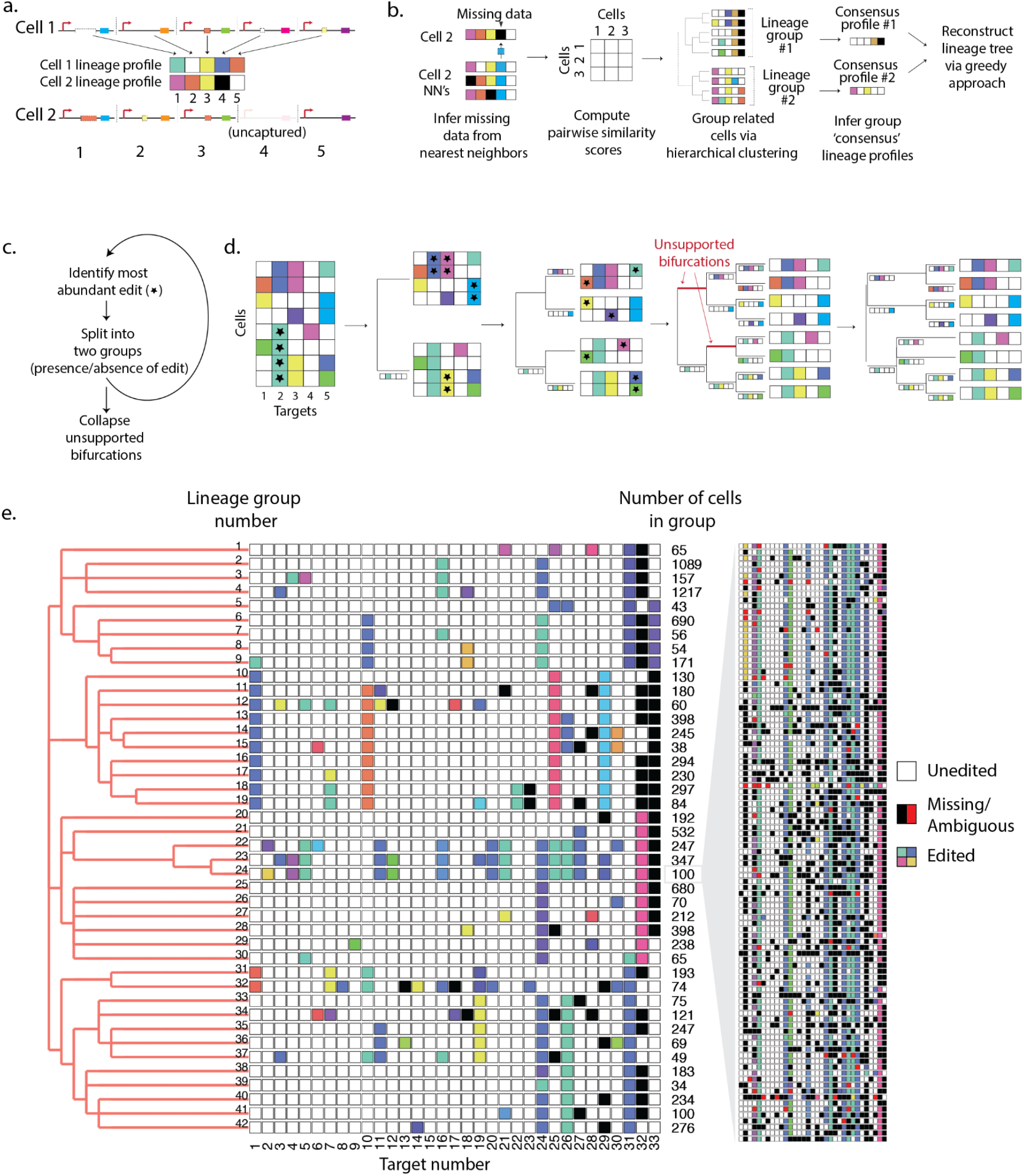
Cell lineage tree reconstruction. (**a**) Visualization of cell lineage profiles. Each unique editing pattern is assigned a unique color. (**b**) Preprocessing of lineage data. Missing data are imputed from nearest neighbors and pairwise similarity scores are computed from corrected lineage profiles. Similarity scores are used to generate a hierarchically clustered tree, grouping related cells. This tree is subdivided into groups of related cells and consensus lineage profiles are generated for each lineage group. The consensus profiles are then used to reconstruct a preliminary cell lineage tree via a greedy approach. (**c,d)** Summary and example of a greedy approach to reconstruct a cell lineage tree. This greedy approach can be performed iteratively on groups of cells within a lineage group to generate a tree with individual cells at the leaves. (**e**) Left: Tree of cell lineage groups (“consensus” editing patterns shown as rows; each column represents a unique target site). Each color represents a unique editing pattern. White: unedited target. Black: targets with missing data for a majority of cells in the group. Number of cells represented by each consensus cell is shown. Inset (right) shows the editing patterns for all 100 cells assigned to lineage group #24. Black: missing targets. Red: ambiguous targets.

The success of this approach is dependent upon two important assumptions: erroneous or missing data is minimal, and convergence events—two or more identical edits occurring independently at a single target site—are rare. We thus set out to optimize the dataset to better fit these assumptions. Sources of erroneous data include PCR and sequencing errors within the target, where a single mismatch in the 70bp (unedited) amplicon would instead appear as a distinct edit. Defining edits is further complicated by the fact that an edit containing both deleted and inserted bases can appear discontinuous when aligned to the reference sequence (*e.g*. see examples within alignments shown in **Figure 2f**). To mitigate errors and misalignments, we required that an edit had to begin within 4 bases of the CRISPR cut site, and that all discontinuous segments be within a maximum of 4 bases from each other (**Methods**). To address missing data, we first defined a similarity metric between cells based on shared edits and used it to identify a set of nearest neighbors for each cell. We then imputed missing and ambiguous edits from these nearest neighbors (**Methods**). Individual cell lineage profiles for a group of closely related cells with missing and ambiguous data shown (black and red boxes, respectively) are plotted in **Figure 3e**.

An additional source of error arises from cross-talk between cellular and target indices during PCR amplification, such that a target sequence derived from one cell becomes associated with the profile of another. A single such error might place a cell far from its true lineage via the algorithm described above. However, although these events are undetectable at the single cell level, they are often obvious when examining groups of closely related cells. To take advantage of this, we sought to pool closely related cells, infer a “consensus” lineage profile for each group (encompassing edits shared by the majority of the group), and generate a preliminary tree of these consensus profiles, such that cells with “contaminating” target sequences would be retained in the group via overall proximity to their neighbors. To identify groups of closely related cells, we again calculated all pairwise similarity scores, and used these as input for hierarchical clustering using Ward’s method. We visually determined the number of clusters into which to subdivide cells, using plots such as the one in **Figure 3e** (right), and computationally inferred a consensus profile for each group. In some cases, where we could explain why an edit did not reach the needed majority for inclusion, automatically inferred consensus profiles were manually corrected (**Methods**). Finally, we applied the algorithm above to the consensus profiles, generating a lineage tree of subgroups of closely related cells.

Since cells within each subgroup contain additional edits beyond the shared edits shown in the “consensus” profile, one can in theory iteratively apply this set of steps to each subgroup, and concatenate the resulting subtrees to derive a single cell-resolved lineage tree. Since our downstream intended application involved comparing pooled expression and chromatin accessibility profiles from groups of closely related cells, and we found that particularly small lineage groups were too noisy for meaningful gene expression and chromatin accessibility analysis, we performed such iterative subdivisions for only a subset of the groups.

For several reasons, we generated an initial tree using only about a quarter of the filtered cells (n = 2,419). First, the hierarchical clustering algorithm used for initial subgrouping has O(n^3^) run time. Second, as described in the previous section, two out of eight batches (1 & 3, **Supplementary Figure 2**) exhibited the most complete lineage profiles, and we reasoned that these would generate the most accurate cell lineage groups into which the remaining cells could be placed via a nearest neighbors approach. Provided that the terminal lineage groups we generate are large enough, we can assume close cell relatives of every cell in the dataset are present within this subset of the overall data. Including all cells, the final tree used for downstream analyses contained 42 lineage groups, ranging in size from 34 to 1217 cells (**Figure 3e**).

This iterative approach of building and concatenating subtrees from root to tip mitigates the probability that recurrent editing patterns at individual targets grossly impact tree structure. For example, if the same edit occurred in two cells independently at target #2, and if one of these events occurred early enough to define an early bifurcation, all descendants of the other cell would be misplaced early during tree reconstruction when employing a greedy approach. However, initial subgrouping of cells based on the full set of edits they contain prevents this problem when at least one of the edits occurs late enough that it does not define the group as part of its “consensus” lineage profile.

Nevertheless, CNAs inferred from expression data occurring over the course of this experiment (discussed in detail in the next section) signaled the presence of two convergence events within lineage data impacting our tree structure. In each case, the convergence events were mediated by a very common editing pattern (**Figure 2f**), and we manually resolved these events to come to the tree structure shown in **Figure 3e** (**Methods**). However, it should be emphasized that with the exception of these two manual changes, the tree shown in **Fig. 3e** was reconstructed solely from lineage profiles, *i.e*. expression data was not used for lineage inference.

### Chromosome copy number alterations inferred from sci-RNA-seq recapitulate the lineage-inferred tree structure

We reasoned that heritable variation in gene expression patterns should visually correlate with tree structure, whereas non-heritable variation should not (**Figure 1a**). To explore this, we aggregated single cell expression profiles within each of the 42 groups described above, and plotted relative group expression as a heatmap (**Figure 4**). Unexpectedly, when genes were arranged by their genomic location, we observed large, continuous stretches of down- or upregulated genes, strong evidence of partial or full chromosomal gain or loss events. HEK293s are pseudotriploid and known to be karyotypically unstable, and an active CRISPR/Cas9 system may also contribute to instability (Y.-C. Lin et al., 2014).

**Figure 4.**
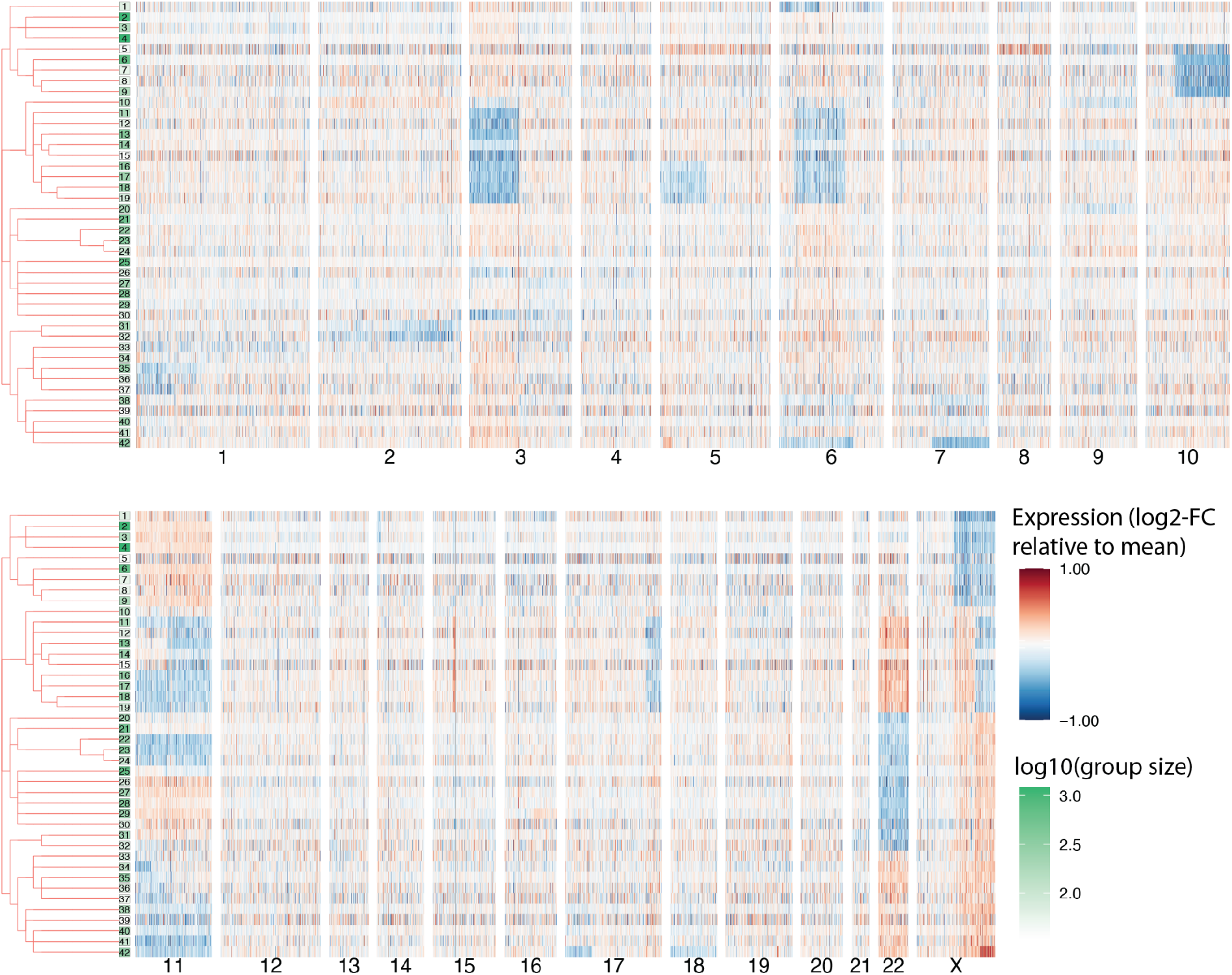
Gene expression in lineage groups arranged by genomic location. Heatmap shows log2-fold gene expression variation relative to the mean expression of each gene across cells. Genes are shown in the order in which they appear along chromosomes in the reference human genome. Log2 fold changes >1 & -1 were manually fixed at these maximum and minimum values for visualization. A minimum mean expression cutoff was applied to remove lowly-expressed genes, leaving 6,241 genes. Green shading of the boxes containing lineage group numbers at the tree leaves is based on the log-scale number of cells per group.

As CNAs are themselves heritable genomic events, we saw an opportunity to use them to validate our CRISPR-inferred tree structure. Strikingly, where present, CNAs were generally concordant with the tree structure inferred from lineage data. In particular, with the exception of full chromosome gains or losses, most CNAs appear to have arisen from a single founder event (**Figure 4**). As described in the previous section and **Methods**, on two occasions, CNAs were used to resolve ambiguity in the lineage data due to convergence events. However, the remaining CNAs shown in **Figure 4** were not used for lineage reconstruction and, importantly, we observed no instances of CNAs contradicting CRISPR-derived lineage relationships.

### Allelic ratios further inform chromosome copy number dynamics across lineages

We next wondered whether we could use lineage-resolved expression data to investigate allele-specific copy number dynamics. Indeed, although we made no direct measurement of copy number, we found that in many cases we could infer copy number based on SNP ratios in sci-RNA-seq data (**Figure 5a**). For example, if a chromosome shows heterozygosity at known SNPs, and we observe allelic ratios of 1:2 across these positions, this chromosome is likely to be present in three copies, while a 1:1 allelic ratio would suggest two or four copies, and a 1:3 allelic ratio would suggest four copies. On the other hand, a paucity of SNPs would suggest regional or chromosome-wide loss-of-heterozygosity, in which case copy number could not be inferred by this method.

**Figure 5.**
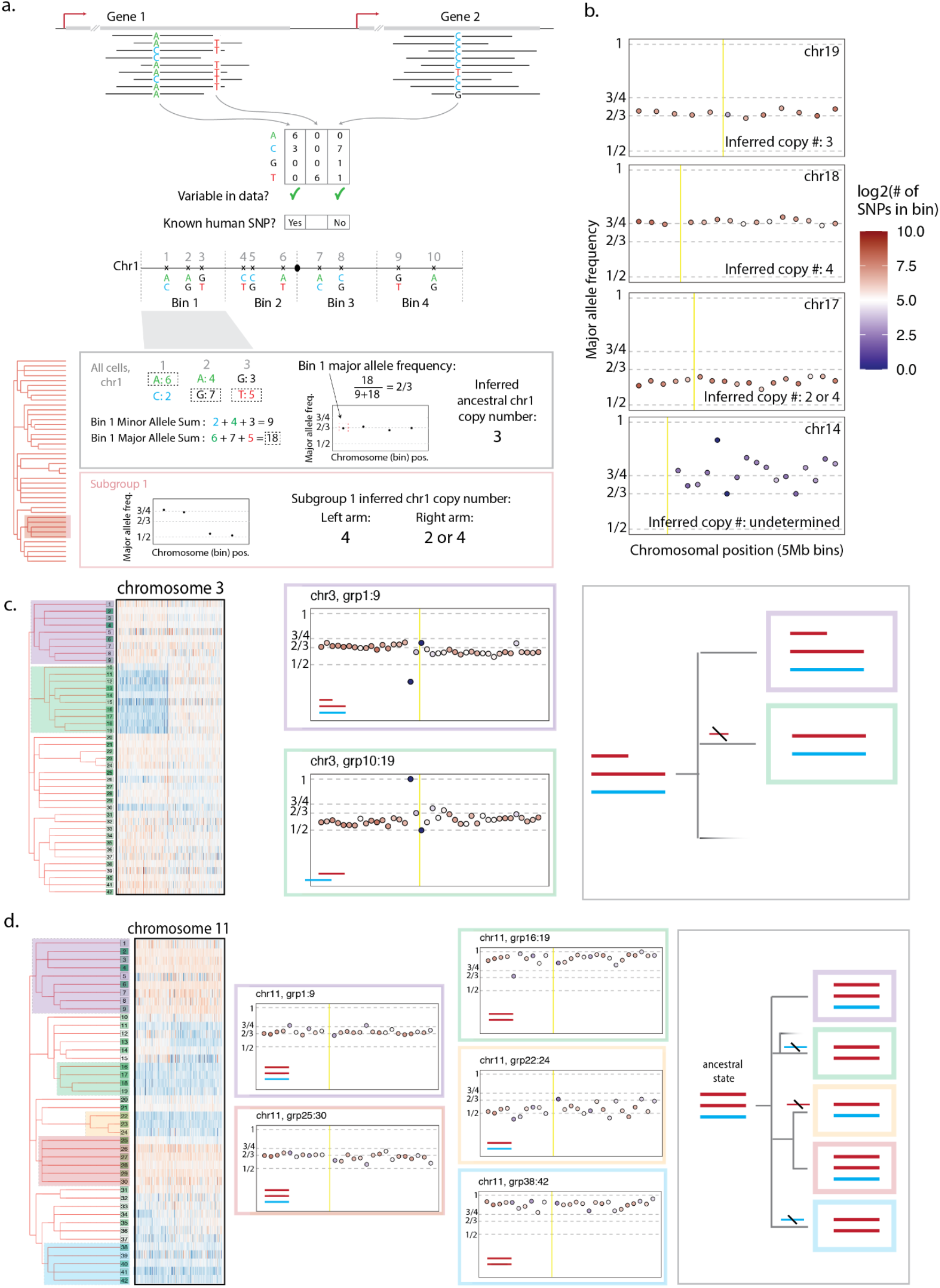
Lineage-resolved allelic ratios inform complex chromosome copy number dynamics. (**a**) A strategy to infer copy number using SNPs from sc-RNA-seq data. First, haplotypic imbalance is assumed and haplotypes are inferred based on base abundance at known SNPs, using all cells. We can then use these to infer the ancestral (or most observed) copy number. Using these haplotypes, we can perform this analysis on subsets of the tree to infer whole or partial chromosome gains or losses. (**b**) Copy number analysis described in panel **a** for chr19, chr18, chr17, & chr14, using all cells. Point fill color represents the number of SNPs found to be heterozygous in that bin, signaling the reliability of this analysis at that location. Yellow line shows the centromere position. (**c**) Subgroup copy number analysis of chr3. Left: expression heatmap as described in **Figure 4**. Middle: Copy number analysis of chr3 for indicated subgroups. Right: schematic of inferred haplotype dynamics. Point fill color represents the number of observed heterozygous SNPs per bin detected when pooling all cells, not just subgroup cells. Yellow line shows the centromere position. (**d**) Subgroup copy number analysis of chr11. Left: expression heatmap as described in Figure 4. Middle: Copy number analysis of chr11 for indicated subgroups. Right: Schematic of inferred haplotype dynamics. Point fill color represents the number of observed heterozygous SNPs per bin detected when pooling all cells, not just subgroup cells. Yellow line shows the centromere position.

We first performed such an analysis on each chromosome using expression data from all cells. Since each genomic position is represented sparsely in sc-RNA-seq data, we divided the genome into 5Mb bins, identified coordinates which appeared to be heterozygous in our data (most frequent base present at in <85% of reads), subsetted these to include only those positions which overlapped known human SNPs (*i.e*. those appearing in dbSNP), and combined counts for SNPs within each 5Mb bin. For this last step, because phasing information was not available, we simply assumed the more abundant alleles at each SNP within a bin were on the same haplotype for binning purposes (as would be expected if homologs existed in unbalanced ratios, at least provided counts are sufficiently high). We then calculated a “major” (most abundant) allele frequency for each bin and plotted these by relative genomic position (**Figure 5a,b**). **Figure 5b** shows several examples of this approach for chromosomes with stable copy number in our dataset, revealing there to be 3 copies of chr19, 4 copies of chr18, and 2 or 4 copies of chr17. Of note, because our heuristic always places the most abundant allele on the same haplotype, we expect a major allele frequency above 1/2 for cases where haplotypes exist in equal copies, *e.g*. as we infer for chr17. On the other hand, chr14 exhibited very low overall heterozygosity at known SNPs together with an unstable ratio, suggesting loss-of-heterozygosity. Consistent with this prediction, the “minor” alleles inferred in chr14 and other chromosomes which exhibit this unstable pattern (**Supplementary Figure 3a**) often do not match known variants founds in the human population, in contrast with inferred minor alleles in chromosomes exhibiting heterozygosity (**Supplementary Figure 3b**). Major allele frequency plots for all chromosomes are shown in **Supplementary Figure 3a**.

We next applied this approach to subgroups of the tree to investigate copy number dynamics during the monoclonal expansion. For example, this analysis revealed a partial loss of an extra copy of the short arm of chr3 impacting only a subgroup of related cells (**Figure 5c**, left panel). Of note, the inferred breakpoint is slightly shifted from the centromere, such that several genes on the short arm are retained. We calculated a binned major allele frequency for the subgroups indicated in **Figure 5c** (left panel), using the major haplotypes we inferred from all cells (**Figure 5c**, right panel). Subgroup copy number analysis (**Figure 5c**, right panel) of groups 1-9 (top, purple) agrees with the predicted ancestral state, whereas the major allele frequency in groups 10-19 has dropped between 1/2 & 2/3 across the whole chromosome. Since heterozygosity appears preserved on the left arm, we infer that the partial chromosome (*i.e*. a copy of the short arm of chr3) was lost in groups 10-19, relative to the ancestral state.

A similar analysis suggested more complex copy number dynamics for chr11, for which multiple full and partial chromosome copy number changes appear to occur at different parts of the lineage (**Figure 5d**, left panel). Performing a subgroup analysis, we observe a pattern consistent with at least three independent full chromosomal losses (**Figure 5d**, middle panel). Intriguingly, these result in different allelic ratios, with loss-of-heterozygosity in two groups (**Figure 5d**, green & blue), and maintained heterozygosity in one (beige). Overall, these analyses highlight the potential of high-resolution, progressive lineage histories to disambiguate copy number alterations, including but not limited to recurrent gains and losses.

### Heritable expression changes unexplained by CNAs are observed throughout the tree

Within genomic regions exhibiting large-scale CNAs, copy number change is the obvious mechanism for differential expression of genes in the impacted region. But other phenomena— *e.g*. epigenetic changes, changes in the levels of upstream regulators, focal CNAs and translocations—might induce heritable expression changes as well. To explore contributions from such sources, we set out to systematically identify examples of heritable expression variation across the tree that were not obviously explained by CNAs.

To this end, we first inferred the boundaries of CNA events between every pair of sister branches (defined as those that share an immediate common ancestor in the tree) using a combination of expression heatmaps (as shown in **Figures 4, 6f**), and pairwise log-fold change plots, where stretches of differential expressed (DE) genes are visible (**Figure 6d**; **Supplementary Figure 5)**.

**Figure 6.**
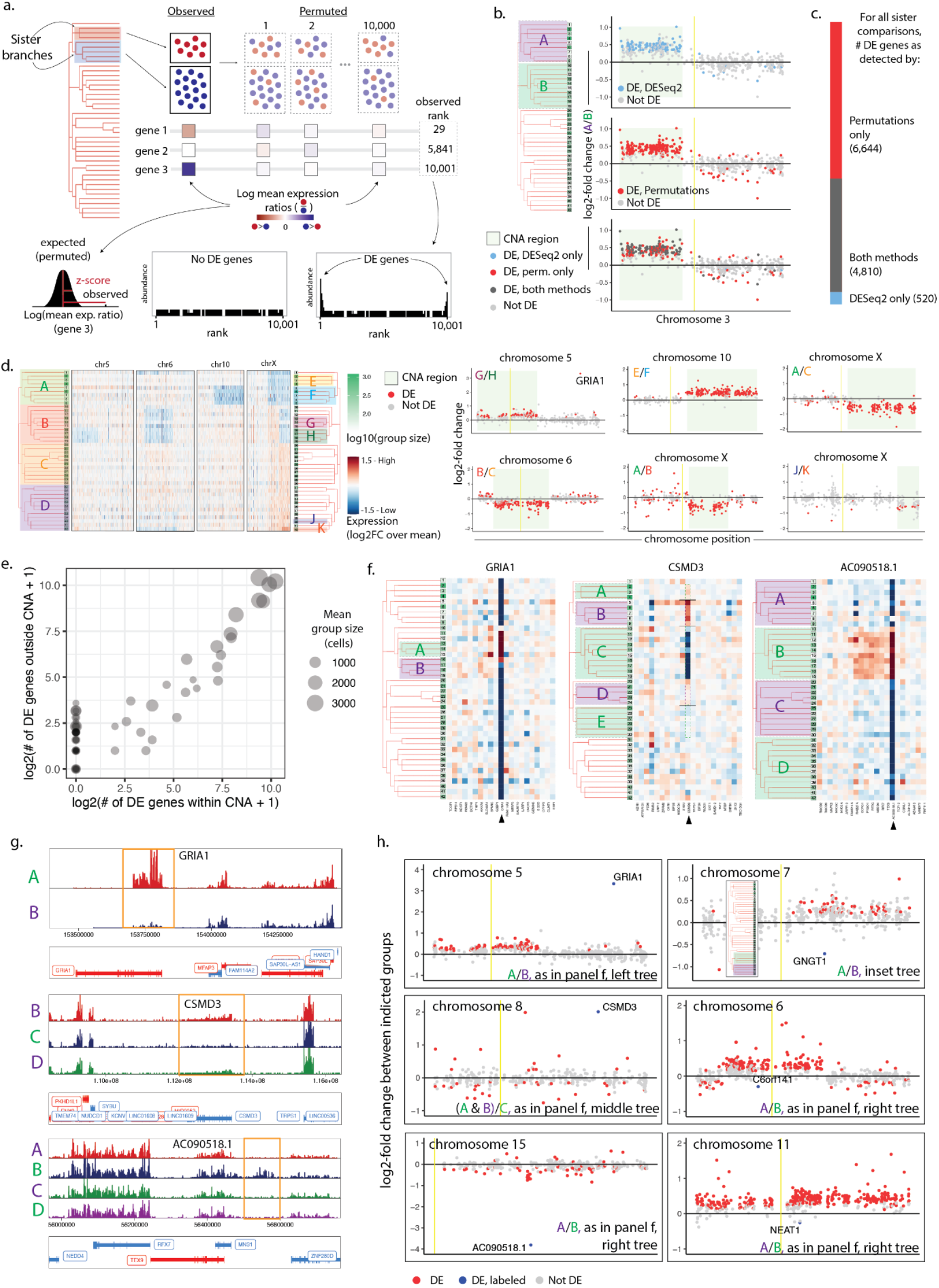
Detecting heritable differential expression within lineage-resolved sci-RNA-seq data. (**a**) A permutation-based strategy for identifying significantly DE genes. (**b**) Comparison of DE genes identified by the permutation method and/or DESeq2, showing log2-fold change expression on chr3 between indicated groups A & B. Yellow bar indicates centromere position. (**c**) Number of DE genes identified using permutations, DESeq2, or both, across every pairwise comparison (66 total) of sister lineage groups (*i.e*. branches sharing an immediate common ancestor in the tree). (**d**) Left: Heatmaps as described in **Figure 4a** depicting CNAs on chrs 5,6,10, & X, with lineage groups indicated on tree. Right: Log2-fold changes of genes on indicated chromosomes between indicated groups, depicting the power to detect DE genes within CNA regions via the permutation approach across groups of different sizes. (**e**) Relationship between the log-scale number of detected DE genes within CNAs and DE gene falling in non-CNA regions per each sister pair comparison. Size of points represents the mean number of cells in the sister pair. (**f**) Heatmaps showing DE expression of *GRIA1, CSMD3, AC090518.1*, and surrounding genes. (**g**) Pileup visualizations of *GRIA1, CSMD3, AC090518.1* in groups indicated on the trees in panel **f**. *AC090518.1* is positioned between *MNS1 & ZNF280D*. (**h**) DE genes showing heritable expression patterns which cannot be explained by detected CNAs. The pair of groups being compared for each plot is indicated on the bottom right, with groups indicated on the trees in panel **f** (except for top-right sub-panel, for which pair of groups is shown in inset tree).

We then sought to evaluate DE between every pair of sister branches, using DE within CNAs as ground truth for sensitivity. Applying DEseq2, which models data as a negative binomial distribution, we observed a substantial number of false negatives—genes within CNAs which were not detected as DE—even between large groups of cells (**Figure 6b**, top panel; **Supplementary Figure 4a**). We thus sought to develop a strategy which would be sensitive to small-magnitude expression changes, while also being robust to large differences in the number of cells between the groups being compared (**Figure 6a**; **Methods**). As a first step, cells from each pair of sister branches are permuted 10,000 times, in each instance creating two groups of the original sizes. For each permuted set, we calculate the log2-fold change for each gene. We then use permuted expression ratios to (a) generate an expected distribution which we can use to calculate a z-score associated with the observed fold change; and (b) rank against the observed expression ratio to assign significance. For a set of genes evaluated for a pair of groups, if none are significantly DE, the distribution of observed ranks is expected to be uniformly distributed; on the other hand, if there are DE genes, we expect to observe their enrichment at the extremes of the rank list. Using an FDR of 5%, we can calculate a set of “significant” ranks (and thus genes) for each pair of groups being compared.

This permutation strategy detected a substantial fraction of genes within CNA regions as differentially expressed (**Figure 6b,c**; **Supplementary Figure 5**). Genes within CNAs across all pairwise comparisons were more likely to be identified by our approach, with lowly-expressed genes within CNAs more likely to be missed by DESeq2 (**Supplementary Figure 4a**; **Supplementary Figure 5**). For example, between groups A & B, 85% of expressed genes (see **Methods** for filtering criteria) within the CNA region on chromosome 3 were identified as DE using our approach, compared with 49% detected by DESeq2 (**Supplementary Figures 4a, 5**). Unless otherwise stated, here we will refer to DE genes as those identified by the permutation approach at an FDR of 5%.

As expected, statistical power decreases with group size, but we nonetheless detected some DE genes within CNAs even between smaller groups (**Figure 6d**; comparisons G/H; J/K). For example, between group J & K (as labeled in **Figure 6d**), containing 234 and 276 cells, respectively, we detect a subset of CNA-associated genes across several chromosomes (**Supplementary Figure 4c**), including *TRIO, SRPK2, & FGF13* (log2-fold changes of -.22, .36, & -.59, respectively). The allelic chromosome copy number analysis presented in **Figure 5** suggests a copy number change from 4 to 5 on chr5 (*TRIO*) & from 3 to 2 on chr7 (*SRPK2*) between these two groups. Since no heterozygosity is observed on chrX, and thus we cannot infer absolute copy number change for *FGF13*.

In total, across 66 pairwise comparisons, we detected 11,454 DE genes using the permutation approach. Of these, 4,810 (42%) were detected using DESeq2, which detected an additional 520 genes not detected by our approach (**Figure 6c**; **Supplementary Figure 5**). Surprisingly, 48% of DE genes detected by permutation analysis could not be directly explained by large-scale CNAs (**Supplementary Figure 5**). The heritable nature of these expression changes may be a product of smaller scale copy number changes, focal genetic or epigenetic differences, or *trans*-effects mediated by heritable events elsewhere in the genome (*e.g*. CNAs or other). Interestingly, when quantified by sister branch pair comparisons, the number of DE genes that we detected outside CNA regions was well correlated with the number of genes within CNAs (Pearson’s r of log-transformed numbers of genes within vs. outside of CNAs = .90, **Figure 6e**), suggesting CNA-mediated expression changes might contribute to heritable gene expression variation through *trans*-acting effects. However, this relationship may largely be explained by the increased statistical power to detect DE genes in larger groups (Pearson’s r of log-transformed number of genes outside of CNAs vs. group size = .76, **Figure 6e**; **Supplementary Figure 4b**).

The most striking heritable expression change which cannot be explained by an obvious CNA was observed in *GRIA1*, a glutamate receptor subunit on chr5 (**Figure 6f-h**, z-score = 28.2, log2 fold-change (FC) = 3.32, between the indicated groups). Markedly elevated expression is observed in lineage groups 11-15 relative to the rest of the tree (with elevated expression in group 16 likely due to misplaced cells). Though we cannot conclusively determine from this data alone whether this expression change is caused by genetic (e.g. focal amplification) or epigenetic factors, it is notable that *GRIA1* is located in a replication transition zone in various cell lines, potentially predisposing it to structural instability (Watanabe et al., 2014). Additional examples of genes exhibiting differential gene expression patterns that track closely with the lineage-derived tree structure appear throughout the tree (**Supplementary Figure 4d**).

Another intriguing example, where multiple expression levels appear to have been stably inherited is observed in *CSMD3*, on chr8 (**Figure 6f-h**). Group B expression is markedly elevated over its sister group A (A/B z-score = -7.30, log2FC = -0.57), while in the branch encompassing both groups A & B, *CSMD3* is even more highly expressed relative to group C (A&B/C z-score = 27.8, log2FC = 2.02). A weaker, but similarly heritable relationship appears between groups D & E (z-score = 3.8, log2FC = 0.34). Such a heritable but labile expression pattern might indicate flexible but relatively stable regulation at this locus. Interestingly, such graded but clone-specific expression patterns were observed with cell type groups in both *Apoe* and *Lmo4* in mouse neurons (Mold et al., 2022). Alternatively, this lability might be explained by local genomic instability. In fact, translocations at a breakpoint near *CSMD3* have been associated with autism in multiple *de novo* cases (Floris et al., 2008), and the *CSMD3* locus is implicated in a wide range of diseases including epilepsy & non-small cell lung carcinoma (Floris et al., 2008; P. Liu et al., 2012; Shimizu et al., 2003). CNAs are particularly common in branch C (**Figure 4**), bolstering the likelihood that a translocation event explains reduced expression in that group.

Even within CNAs, we observe single gene expression changes which deviate strongly from the expected copy number ratios. An intriguing example is the transcript *AC090518.1*, which normally exhibits testis-specific expression, and is located within a short stretch of genes with modestly elevated expression on chr15 consistent with a CNA (**Figure 6g,h;** *AC090518.1* is located between *MNS1 & ZNF280D*). This transcript’s markedly increased expression well beyond that of its neighbors (log2-fold change (A/B) = -3.82, A/B z-score = -28.67), points to a possible translocation (or tandem duplication) event, exposing it to a new regulatory context. Chromosomal rearrangements are a hallmark of cancer progression, and tracking such small-scale events may reveal the mechanism behind biologically-meaningful expression changes. The genes *GNGT1, C6orf14*, and *NEAT1*, all lie within CNA regions but show heritable expression changes in the opposite direction of surrounding genes (z-scores -7.40, -4.10, -8.84, respectively, **Figure 6h**). Such patterns may indicate expression compensation or selection for particular expression levels. In fact, both *GNGT1 & C6orf141* have been associated with cancer prognosis (Yang et al., 2019; J.-J. Zhang et al., 2021), with *C6orf141* playing a direct role in cell proliferation. *GNGT1* was designated a hub gene in non-small-cell lung cancer, suggesting its misexpression may have widespread downstream consequences which would also appear heritable. *NEAT1*, a long non-coding RNA with a known epigenetic role in a variety of cell types, may also stably modify expression of multiple downstream target genes (Wang et al., 2020).

Here, lineage relationships enabled us to identify stably-inherited expression changes which may not otherwise be obvious among non-heritable expression fluctuations. In most cases, however, it is not possible with this data alone to determine the mechanistic basis for this differential gene expression (*e.g. cis-*genetic, *trans*-genetic vs. epigenetic). We next sought to distinguish between these possibilities by additionally tethering chromatin accessibility information to this same lineage tree.

### Collecting lineage information alongside single cell chromatin accessibility profiles enables tethering of gene expression and chromatin accessibility

Both genetic and epigenetic phenomena can potentially underlie what we observe as heritable expression changes, and measuring expression alone is often not sufficient to disentangle these from one another. Coassays of single cell expression and chromatin accessibility may provide more insight, but contemporary methods result in relatively sparse profiling in any given cell. However, since heritable states are presumably shared by cells with similar lineage histories, we can theoretically measure these features independently in clonally related cells and link them retrospectively based on lineage relationships (**Figure 7b**). Furthermore, pooling of single cell chromatin accessibility profiles of closely related cells, as we did with expression profiles, increases the power to detect changes. To this end, we developed a method to capture lineage-associated transcripts alongside sci-ATAC-seq (Cusanovich et al., 2015, 2018), *i.e*. to concurrently profile single cell lineage relationships and chromatin accessibility states (**Figure 7a**). sci-ATAC-seq is a pool-split approach where genetic material undergoes two rounds of molecular indexing, such that DNA from each cell is ultimately associated with a unique pair of indexes. To associate lineage information with sci-ATAC-seq profiles, we devised a strategy to concurrently index mRNA transcripts containing recorded lineage information at each sci-ATAC-seq indexing round, via reverse transcription and PCR, such that both features can be retroactively linked to a single cell via index combinations (**Figure 7a**; **Methods**).

**Figure 7.**
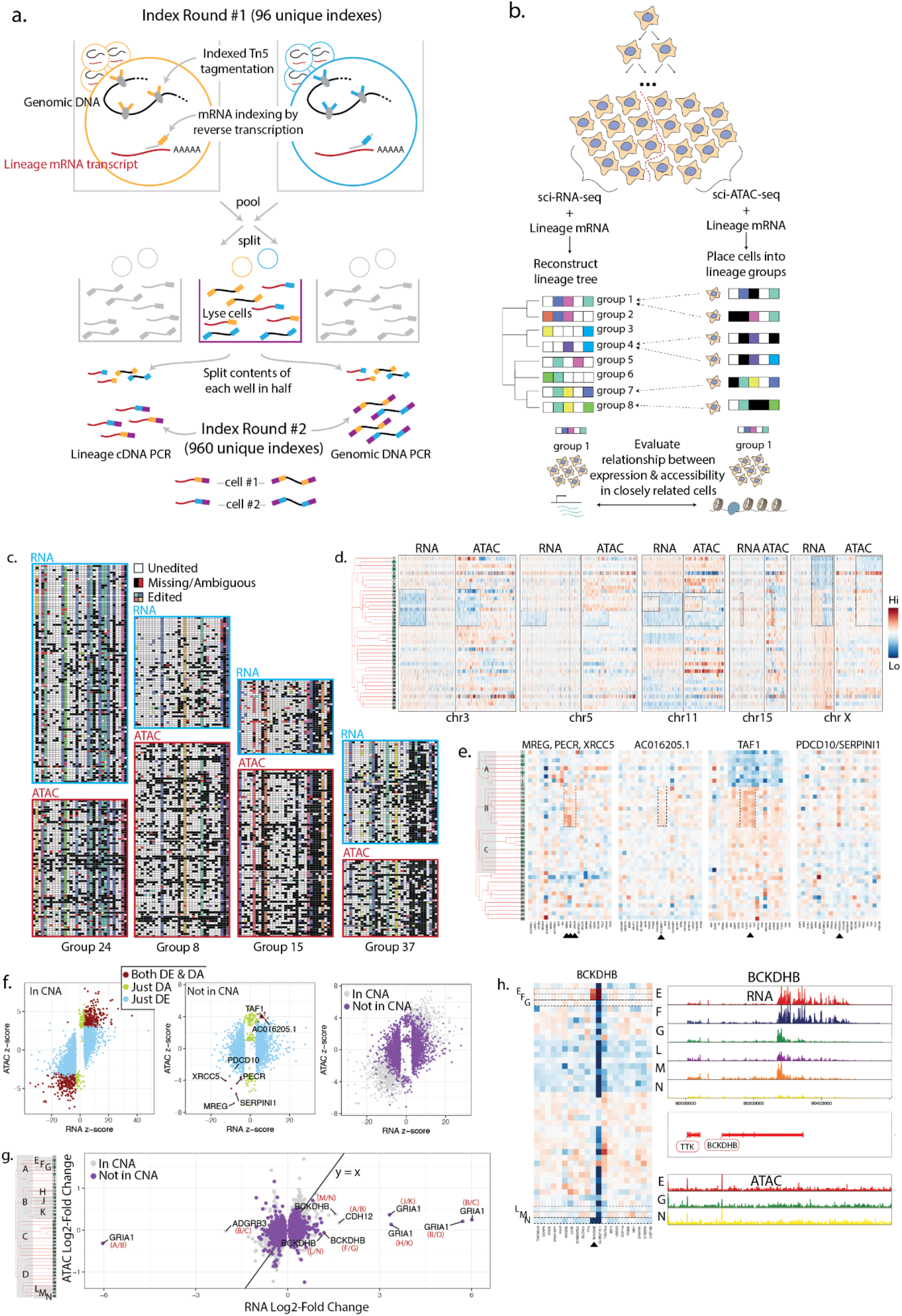
Collecting chromatin accessibility data (via sci-ATAC-seq) alongside lineage profiling, and evaluating its relationship to expression in closely related cells. (**a**) A combinatorial indexing strategy to concurrently capture chromatin accessibility and lineage mRNA from the same single cell. (**b**) Schematic depicting how expression (sci-RNA-seq) and accessibility (sci-ATAC-seq) are linked via lineage information. Lineage-traced cells are split in half, and lineage profiles are captured separately alongside each single cell feature. A lineage tree was reconstructed from cells with concurrently profiled expression, and lineage profiles of cells with concurrent accessibility profiling were used to place cells into previously defined lineage groups via nearest neighbors. The relationship between expression and accessibility of closely related cells could then be evaluated. (**c**) Lineage profiles of individual cells within four clonally related groups collected alongside either sci-RNA-seq or sci-ATAC-seq. (**d**) Heatmaps showing the relative expression (RNA) and accessibility (ATAC) across the 42 lineage groups, calculated for each gene (RNA), and for each 1MB bin (ATAC) for five selected chromosomes. Genes & bins are ordered by their chromosomal position. Dashed boxes indicate chromosomal regions with visually consistent copy number changes across the tree. (**e**) Heatmaps showing relative expression for a subset of genes which are both DE and DA, and including 10 positionally adjacent genes on either side. Associated RNA & ATAC read pileups are shown in **Supplementary Figure 5d**. (**f**) Left: Relationship between expression and accessibility changes evaluated within gene bodies plus 5kb upstream of the TSS, calculated using the permutation approach described in **Figure 5a**. Only genes within CNAs are shown. Each point represents an expression/accessibility change at a single gene for a pair of sister lineage groups (and thus a gene may be represented more than once). Points are colored by their DE and DA status. Middle: Analogous to the left plot, except including only genes *outside* of CNAs. Labeled genes are referenced in the text. Right: Overlay of left and middle plots. 10 outlier genes, where noise was likely due to low expression/accessibility, were removed from the middle and right plots. (**g**) Relationship between RNA and ATAC log2-fold change (as opposed to z-score). Each point represents an expression/accessibility change at a single gene for a pair of sister lineage groups (and thus a gene may be represented more than once). Outliers discussed in the text are labeled with gene name and pair of sister groups as indicated on the tree. Because small groups result in noisy data, comparisons involving at least one small group (<100 cells) were removed. An expression cut-off was also applied to reduce visual noise, leaving the 45% of comparisons with the highest expression. (**h**) Left: Heatmap of relative expression of *BCKDHB* and surrounding genes, respectively. Right: Pileup of expression and chromatin accessibility data for the indicated groups (as labeled on tree in panel **g**). Log2-fold change between groups F&G: 1.18 (RNA), -0.06 (ATAC); groups L&N: 1.13 (RNA), - 0.12(ATAC).

We applied this method to the remaining cells from the lineage/expression capture experiment, and filtered cells to those for which we collected both chromatin accessibility profiles and suitable lineage information. Since a lineage tree has already been built, lineage profiles captured alongside sci-ATAC-seq need only be complete enough to accurately place them into existing lineage groups. Keeping cells with at least 5 captured targets of which at least one was edited, with more unambiguous than ambiguous editing events (the latter likely representing doublets), we retained 12160 cells with lineage information. In this group of cells, a median of 20 unique targets were captured per cell (**Supplementary Figure 5b**). We next filtered on chromatin accessibility profiles. Chromatin fragment lengths exhibited the expected nucleosomal peaks (**Supplementary Figure 5a**), and filtering on UMI counts yielded a total of 9014 cells (median non-mitochondrial UMI count: 1601; mean UMI count: 6491; minimum 32 UMIs, **Supplementary Figure 5a**).

To place these cells into existing clonal groups, we first computed a weighted similarity score based on lineage profiles for each ATAC-associated cell with each RNA-associated cell. We then placed cells into existing groups based on nearest neighbors (**Figure 7b**). Encouragingly, the relative group sizes of ATAC-associated cells correlated well with the original group sizes (**Supplementary Figure 5c**). Moreover, lineage profiles collected alongside accessibility were visually consistent with those collected alongside expression within tethered groups (**Figure 7c**). Together, these data suggest that cells were accurately placed into lineage groups, and thus we can expect analogous heritable states to be reflected in expression and accessibility measurements within a group.

### Using lineage-tethered chromatin accessibility and expression profiles to investigate mechanism of heritable expression

Although sci-ATAC-seq is primarily used to measure local chromatin accessibility changes, copy number changes should also be apparent since they affect the amount of DNA available for tagmentation. Thus, if paired expression and accessibility measurements truly capture closely related cells, CNAs observed in expression data should also appear in accessibility data. To visually evaluate CNA concordance, we quantified relative sci-ATAC-seq read counts across 1MB windows of the genome for each lineage group and generated heatmaps analogous to those shown in **Figure 4**. Indeed, we observed striking agreement in CNA patterns between expression and accessibility data (**Figure 7d**), further confirming lineage profiles do link close cell relatives. To determine if CNAs were measurable in accessibility data at the gene level, we evaluated accessibility within gene bodies, including 5kb upstream of the TSS, once again using the permutation strategy described in **Figure 5a**. We found that within CNAs, RNA and ATAC z-scores are strongly correlated at genes which are DE, DA, or both (Pearson’s r = .73, **Figure 7f**, left panel), while much more limited correlation is observed outside of CNA regions (Pearson’s r = .16, **Figure 7f**, right panels).

Since copy number differences are often observable at the gene level in ATAC data, we wondered if we could use gene body accessibility outside of large CNAs to identify genes whose DE status is likely due to small genomic amplifications or deletions, affecting one or a few genes. Correlated DE and DA status may alternatively indicate a regulatory change, but such DA is more likely to be promoter-specific; in this case, we would expect a higher promoter-specific signal, while evaluating DA across the whole gene body could dampen such localized signal (Nair et al., 2021). 21 genes outside of CNAs are both DE and DA (**Figure 7f**, middle panel), making them good candidates for residing in short CNAs. In fact, three of these—*MREG, PECR, XRCC5*—are adjacent genes on chr2, with higher expression in group B relative to group A, despite similar expression outside of this region (**Figure 7e**; **Supplementary Figure 6d**). This pattern strongly suggests that a focal amplification occurred at this locus, explaining the increase in transcript abundance. Similarly, *AC016205.1* on chr18 & *TAF1* on chrX are both DE and DA between the groups indicated in **Figure 7e**, and also appear within short stretches of genes with elevated expression. A pileup of ATAC data, showing the positions of Tn5 insertions across *TAF1*, shows elevated signal across the whole gene body as well as the neighboring gene *OGT*, validating our prediction. A small CNA is also likely on chr3, where elevated expression is observed in DA gene *SERPINI1* and nearby *PDCD10* (**Figure 7e**; **Supplementary Figure 6d**, *SERPINI1* does not appear on the heatmap due to low expression level.). Although *PDCD10* is not significantly DA by our metrics, it lies in the vicinity of genes which are (**Figure 7f**, middle panel). Pileup of ATAC reads in this region supports this prediction, with denser coverage of reads across the gene body of *SERPINI1* in group B (**Supplementary Figure 6d**, right panel). These data suggest that paired expression and accessibility data can help identify small copy number changes.

We next sought to use accessibility data to identify genes whose expression changes are <u>unlikely</u> to be mediated by copy number changes. If a heritable expression change is triggered by a simple gene copy number change, we expect a linear fold-change concordance between expression and gene body accessibility. If, on the other hand, an expression change is due to other factors, such as abundance of an upstream regulator or change in its regulatory context, these features are not necessarily expected to be linearly correlated. Though log2-fold changes at single genes between variable size groups are inherently noisy, especially in ATAC data, outlier DE genes are especially likely candidates for non-copy number mediated heritable states. We thus further inspected several such outliers, where expression change greatly exceeds accessibility change (**Figure 7g**).

Between groups A & B as indicated in **Figure 7g**, the expression change in *GRIA1* is 19 times greater than its gene body accessibility change (RNA log2-fold change = -6.04; ATAC log2-fold change = -.32), suggesting genomic amplification is very unlikely to be the cause of this expression change. Similarly, the expression changes observed in *BCKDHB, CDH12, and ADGRB3* (**Figures 7h, Supplementary Figure 5e,f**) between the indicated groups greatly exceed gene body accessibility changes (log2-fold change shown in figure or legend). The absence of significant accessibility change in *BCKDHB* in particular allows us to rule out a focal amplification of the 3’ end of the gene as an explanation for high RNA read coverage specifically in that region in groups E & F (**Figure 7h**). A more likely explanation is that a different transcription termination site was used.

Beyond copy number changes, heritable changes in accessibility at regulatory regions would signal an epigenetic origin to expression variation. We thus identified peaks in ATAC data, both in the entire dataset as well as in lineage-specific subgroups internal to the tree, and looked for DA peaks within 5kb of TSSs or within the gene body between every pair of sister groups near genes found to be DE. We did not observe any DA peaks in these regions. Consistent with this, Kiani *et al*. recently showed that accessibility and expression changes are not well correlated in single gene perturbation experiments (Kiani et al., 2022). Others have observed a similar lack of concordance between accessibility changes and expression level (Hota et al., 2020; Y. Zhang et al., 2020).

Together, these data illustrate the potential of lineage-based coupling of expression and accessibility data to help distinguish between potential mechanistic explanations for heritable expression changes.

## Discussion

Here, we have shown how tethering single cell expression and chromatin accessibility profiles via lineage relationships facilitates the detection and characterization of heritable gene expression changes. Surprisingly, even in a non-differentiating cell line, we observed abundant, progressively-acquired heritable expression changes. Some differentially expressed genes had an obvious genetic origin—copy number changes impacting multiple adjacent genes, while many others showed stable, lineage-associated expression but with less clear origins. The explanations for this later category might include epigenetic changes within nearby regulatory sites, changes in abundance of upstream regulators, the acquisition of new regulatory contexts via genomic rearrangements, and/or focal genetic changes, amplifications, or deletions. Above, we have shown that our approach of profiling multiple features in closely related cells can, at least in some cases, be used to distinguish between these possibilities.

Clonal tracking, achieved via various methods across diverse systems, has revealed the presence of biologically important heritable states. For example, combining Luria-Delbrück fluctuation analysis with RNA-seq, Shaffer *et al*. found rare, but clonally stable expression states which predisposed cancer cells to drug resistance (Shaffer et al., 2020). Intriguingly, these states were in some cases reversible, suggesting an epigenetic origin. Goyal *et al*. confirmed the presence of clone-specific responses of cancer cells to various drug treatments using a clonal barcoding approach (FateMap) (Goyal et al., 2021). Mold *et al*. made use of ‘natural’ clonal barcodes—T-cell receptors in lymphocytes—and found that clonal lymphocytes responded more similarly to vaccination than more distantly related cells (Mold et al., 2022). Using an *in vivo* transgenic barcoding strategy (TREX, (Ratz et al., 2021)), they found that in mouse neurons, gene expression states mimicked clonal structure, even among different clones of the same cell type. Finally, He *et al*. investigated the timing of cell fate restriction in organoids with iTracer, a system which includes an initial and an induced round of clonal barcoding (He et al., 2021). These studies present intriguing examples of heritable expression but are limited in terms of fully distinguishing between potential underlying causes.

We envision that THE LORAX may be applied to such systems, enhancing our ability to detect heritable events and explain their mechanistic origins. First, progressive lineage labeling increases the likelihood of detecting rare heritable events, as finer-scale, temporally-resolved clonal labeling produces more homogenous clones. Progressive labeling may be particularly useful for detecting events which are stable over multiple cell divisions but reversible, since both the acquisition and reversal may be captured via a finely-tuned lineage recording system. Second, the addition of a chromatin accessibility measurement alongside clonal labels may help resolve the mechanisms behind clonal expression stability. Genetically-mediated expression variation is likely during cancer progression, where copy number changes ((Harbers et al., 2021), loss of heterozygosity (Nichols et al., 2020), and chromothripsis (Cortés-Ciriano et al., 2020 — widespread fragmentatio|n and reassembly of genetic material—are commonly observed. We have shown above that such events may be inferred using our approach. On the other hand, myriad epigenetic changes accompany cell fate commitment during organoid and organism development, and concurrent lineage tracing and RNA and ATAC profiling in closely related cells may illuminate the order of events which give rise to progressive cell type divergence (Thomas et al., 2011). In these systems and others where cell state diversification is taking place, it is likely that lineage-resolved ATAC-seq will show clone-specific enhancer and promoter accessibility changes beyond what we observed here, which may explain heritable expression variation. In fact, profiling clonal T-cell populations expanded *in vitro* using bulk ATAC- and RNA-seq, Mold *et al*. found clone-specific accessibility changes at regulatory regions, with enrichment near clonally differentially expressed genes (Mold et al., 2022).

Our work presents some advances in CRISPR-based lineage tracing, and also highlights some fresh challenges. First, encoding lineage at many independently-integrated loci rather than at tandem loci expressed as a single transcript eliminates the chance that a large deletion removes neighboring CRISPR targets, supports larger deletions, and enables efficient capture of larger insertions. These features in turn reduce both the rate of missing lineage information and the probability of convergence events. Second, we show that NN-based inference of missing data in individual cells and subsequent pooling of cells to generate “consensus” profiles prior to lineage reconstruction (and iteratively generating subtrees from these consensus groups) reduces the likelihood of misplaced cells early in the reconstruction process. Though we demonstrate the usefulness of this approach when a “greedy” algorithm is used for reconstruction, it is applicable even to methods which primarily use traditional phylogenetic reconstruction approaches (*e.g*. maximum likelihood) (Gong et al., 2021; Jones et al., 2020; Konno et al., 2022), since the sheer number of cells often makes early “greedy” subgrouping necessary. Finally, these lineage recording and analysis approaches are compatible with other recent advances in lineage recording technology, like DNA Ticker Tape (J. Choi et al., 2021), where successive insertions as a single locus greatly simplify ordering of lineage-encoding events. Integrating multiple such loci would enable higher resolution trees, and the approaches presented here can be used to order events occurring at distinct recording loci, where event ordering is not so straightforward.

Our work also highlights some unresolved challenges within the CRISPR-based lineage tracing field. First, fine control of editing rate remains elusive; we observed abundant editing in some lineages, while most targets in others remained unused. Loss or silencing of the Cas9-expressing genomic locus might explain lineage-specific reductions in editing efficiency, while position effect variegation in cutting or editing rates might explain variation in usage or recovery across targets. Second, though we observed a great diversity of editing patterns, they are not evenly distributed, with the top three edits frequently occurring independently. This phenomenon can in part be addressed with careful target design to avoid regions of microhomology (W. Chen et al., 2019; Sfeir & Symington, 2015). Third, though the design of our construct allows for large indels relative to other methods, relying on double strand break repair for editing diversity still presents a risk that a recorded event will not be reliably captured due to indel size. Finally, frequent DSBs (which may themselves be contributing to the CNAs observed here), and the persistently high expression of transgenes (which are prone to silencing) may not be compatible with organismal or ES cell-derived systems. Excitingly, these challenges are addressed in large part by DNA Ticker Tape, which leverages prime editing to introduce diverse insertional edits to a target site in an ordered manner, without requiring double-stranded breaks (J. Choi et al., 2021).

The logical core of THE LORAX–pooling cells based on genetically-encoded labels captured alongside multiple genomic and/or epigenetic features to evaluate the relationship between those features–is broadly applicable to any system amenable to genetic barcoding. Systems where static barcodes (*e.g*. CellTag (Guo et al., 2019)) are used to interrogate clone-specific heterogeneity, are an obvious candidate, but labels need not necessarily mark clonal populations. For example, sgRNAs in CRISPR perturbation screens can be used to tether multiple single cell molecular measurements. Importantly, combinatorial indexing approaches are not required here, as both short barcode integrants and sgRNAs can now be captured alongside scRNA-seq (Biddy et al., 2018; Dixit et al., 2016; Guo et al., 2019; Rodriguez-Fraticelli et al., 2018; Weinreb et al., 2020) and scATAC-seq (Pierce et al., 2021; Replogle et al., 2020; Rubin et al., 2019) via droplet-based methods.

In some applications, THE LORAX has several advantages over traditional co-assays of expression and accessibility where both features are measures in the same single cells (Cao et al., 2018; S. Chen et al., 2019; L. Liu et al., 2019; Ma et al., 2020; Xing et al., 2020; Zhu et al., 2019), as well as computational integration methods which merge single cell expression and accessibility datasets (Y. Lin et al., 2021; Stuart et al., 2019). First, existing co-assay methods are relatively low resolution compared with methods which profile each feature separately; thus, associating single-feature profiles via lineage relationships improves resolution at the single cell level. Second, by aggregating profiles of closely related cells, we achieve higher statistical power to detect even rare, heritable events. Third, though computational integration is possible in datasets composed of a variety of cell *types*, it is less feasible in ones composed of different cell *states* where well-separated clusters are not expected and stochastic factors often drive within-cluster positioning. THE LORAX enables overlaying of expression and accessibility datasets without making *a priori* assumptions about their relationship, as is necessary during computational integration.

In summary, we have shown that (a) progressive recording of lineage information across distinct genomic loci, and their high rate of recovery alongside sci-RNA-seq, enables accurate reconstruction of cell lineage trees; (b) aggregating expression profiles of closely related cells reveals abundant, and progressively acquired heritable expression variation, even in non-differentiating cells; and (c) we can investigate the relationship between multiple features—like expression and chromatin accessibility—by tethering them via concurrently captured lineage profiles.

## Materials and Methods

### CRISPR lentiviral target construct & Cas9 construct generation

Target/sgRNA construct: In order to integrate CRISPR targets and sgRNAs into the genome, we modified the CROPseq vector (Datlinger et al., 2017) (Addgene ID 86708), which expresses an sgRNA and a PolII transcript. We integrated a CRISPR target construct after the WPRE, such that it is expressed off the PolII promoter (sequence and location shown below). Target constructs were identical except for a unique 10bp barcode. sgRNAs matched the targets and were thus identical across all uniquely-barcoded constructs. A primer binding site was placed 35bp upstream of the CRISPR cut site, such that the target accommodates a 70bp deletion. The sequencing and computational processing scheme enables capture of insertions of >105 bp. (see <u>Computational processing and edit calling from lineage target sequencing data</u>))

Target insert:

TCCAAGCTCCATAGGTCCAACTCAAGCTTAGTTCCTATACTGATTCCAAGCCATGGTACCATAGCAGATGATCCATTTAGAGCCTGGCTGGTCTCCTGGGAGGTCAACCTTGGAGACTAAGACCTTACGNNNNNNNNNN

Unique target barcode

gRNA binding site

Forward primer binding site

Position of insertion after WPRE between sequences shown:

TCCCCGCGTCGACTT[INSERTION SITE]TAAGACCAATGACTT

Primer binding sites:

Forward: CTGATTCCAAGCCATGGTAC

Reverse: GACTTACAAGGCAGCTGTAG

A modified version of the doxycycline-inducible SpCas9 lentiviral plasmid (https://www.addgene.org/50661/) was used in this experiment. This construct contains an auxin inducible mAID sequence (cloned from pMK288 (mAID-Bsr), Plasmid #72826, Addgene) This degron sequence was not used in this experiment. Doxycycline was not used to induce this construct -- instead, we relied on known leaky expression to achieve a low level of editing. The full construct sequence is available on <u>Benchling</u>.

### Cell line generation

HEK293 (ATCC, CRL-1573) were first transduced with the barcoded target/sgRNA modified CROPseq vector at high MOI and single cells were sorted to grow clonal populations. Targets were counted by PCR amplifying and sequencing the unique barcodes. A clone containing 31 unique barcodes was chosen.

To induce editing, cells were transduced with the doxycycline-inducible Cas9 lentiviral construct described above, selected for Cas9 integration using Blastocidin, and single cell sorted such that all profiled cells arose from a single founder cell. The Cas9 construct was not induced with doxycycline; instead, we made use of its known propensity for leaky expression without induction to produce slow editing. After 35 days in culture (DMEM), passaged every 2-3 days using trypsin, editing efficiency was evaluated by bulk PCR of the target regions, and a single clonal edited population was chosen for further exploration. A portion of the resulting cells were collected and processed immediately in a target+sci-RNA-seq capture experiment, and a portion was frozen in liquid nitrogen for later target+sci-ATAC-seq processing.

### Capture of CRISPR targets alongside sci-RNA-seq

The sci-RNA-seq 2-level protocol for methanol-fixed cells described in Cao *et al*. 2017 (Cao et al., 2017) was modified to concurrently capture CRISPR target mRNAs. A single 96 well plate was used for the first round of indexing, and 8 96-well plates were used in the second round, with 25 cells sorted into each well.

The following modifications were made:

1. To index the lineage target mRNA during the first round of indexing, we added a 1um of 10uM indexed target-specific reverse transcription primer in addition to the oligo-dT primers. Reverse transcription primer sequence: ACGACGCTCTTCCGATCTNNNNNNNNTTGGTAGTCG ctacagctgccttgtaagtc UMI RT index (well-specific sequence)
2. After Tn5 tagmentation, lysis, and ampure bead purification, cDNA was eluted in 10ul of buffer EB (Qiagen). Then half of the contents of each well were transferred to a second 96 well plate. In one plate, PCR and sequencing of the transcriptome was performed as described. The other plate was used for amplification of the lineage targets, with well-specific primers indexed to match well-specific transcriptome indices. Lineage targets were PCR amplified using the KAPA HiFi HotStart ReadyMix (Roche, KK2602) with primer sequences below and elongation time of 1 minute and an annealing temperature of 65°C. All other steps were consistent with the KAPA protocol provided by manufacturer. PCR primers: Forward (unindexed): CAAGCAGAAGACGGCATACGAGATTTGGTAGTCGGTGACTGGAGTTCAGACGTGTGCTCTTCCGATCTCTGATTCCAAGCCATGGTAC Reverse (indexed): AATGATACGGCGACCACCGAGATCTACACTTCTACCTCAACACTCTTTCCCTACACGACGC TCTTCCGATCT PCR index (well-specific sequence) PCR index (plate-specific sequence) After PCR, all wells were pooled and a 0.8x AMPureXP bead cleanup was performed prior to sequencing.
3. Paired-end sequencing of the lineage target PCR products was performed using a 300bp Illumina sequencing kit (Miseq), with 148 bases sequences from each end (along with standard 10bp index reads, which are associated with the second round of indexing). The first index as well as the UMI appear in R1 and are parsed during downstream computational processing. 10% PhiX was added for sequencing to address sequence homogeneity.

### Capture of CRISPR targets alongside sci-ATAC-seq

The concurrent lineage target + chromatin accessibility capture protocol builds upon the 2-level sci-ATAC-seq protocol presented in Cusanovich *et al*. (2015) (Cusanovich et al., 2015). The following modifications were made:

1. Lysis buffer was supplemented with SuperaseIN (ThermoFisher AM2694).
2. Reverse transcription of lineage target mRNA:For s first round of lineage target indexing, reverse transcription was performed prior to tagmentation in the first set of wells. After lysis, 5000 nuclei (2ul) were distributed per well of a 96 well plate, along with reagents for the first step of reverse transcription: 0.25ul dNTPs (10mM) & 1ul of indexed the reverse transcription primer described above (at 2uM). The plate was then incubated at 55C for 5 minutes, and immediately chilled on ice. Reagents from the SuperScriptIV (ThermoFisher, 18090010) kit were then added to each well (1ul buffer, .25ul DTT, .25ul SSIV enzyme, .25ul RNAseOUT (ThermoFisher, 10777019). The plate was then incubated at 55C for 10 minutes, and immediately chilled on ice.
3. Buffer exchange following reverse transcription: 60ul of nuclei lysis buffer was added to each well. Nuclei were then pelleted by centrifugation at 300g for 5 minutes in 4°C. 57ul were then carefully removed from each well, taking care not to disturb the pellet.
4. After sorting nuclei (25 nuclei per well) into a solution containing SDS & inclubating to insure Tn5 inactivation and lysis, the contents of each well are split in half across two plates. One plate underwent indexed DNA PCR amplification in accordance with the sci-ATAC-seq protocol; the other underwent a 2X AmpureXP bead purification to remove SDS, followed by PCR amplification as described above. Primer cleanup and sequencing of lineage target amplicons was performed as described above.

### Initial computational processing of sci-RNA-seq data

Sequencing was performed as previously described (Cao et al., 2017). Reads were adapter-trimmed using trim_galore and aligned to the reference genome (hg38) using STAR. Non-unique mappers were removed. Reads were then deduplicated using a custom script (190223_sciRNA_remove_duplicates.cpp), taking into account both UMIs and cell indices to call a duplicated read. Only cells with at least 2048 deduplicated non-mitochondrial UMIs were used for subsequent analyses.

A custom script (190704_process_sciRNA_mapped_file.cpp) was used to map reads to genes. Reads which overlapped multiple genes but only fell in an exon in one gene were counted towards that gene.

RNA processing to generate the cell by gene raw counts file is implemented in script 190807_sciRNA_wrapper_ALL.txt, with user-defined UMI cutoff of 2^11.

### Computational processing and edit calling from lineage target sequencing data

Targets were enriched from the cDNA as described above and sequences on the Illumina Nextseq or Miseq 300 cycle kit, with paired end sequencing. Read pairs (150b from each end on Miseq; 148 from each end on Nextseq) were merged using PEAR. Since large insertions can possibly result in pairs which do not overlap, we took reads which were unable to be merged and looked for features (common sequence near barcode, primer binding sites) which indicated reads from the correct location. We then pasted the pairs into a single read, and used the combined insertion sequence in our analysis. Thus, insertions of >105bp could be captured, as long as the amplicon could cluster efficiently on the sequencer chip.

Merged reads contain UMIs (first 8bp), reverse transcription index (index #1 of combinatorial indexing - next 10bp), and a target ID (obtained by searching for flanking sequences). These features were first extracted from the reads (191203_CROPt_make_UMI_RT_BC_seq_output_file.cpp, within wrapper script 191203_CROPt_Step2_collapse_UMIs_wrapper.txt), and the remaining sequences were collapsed by UMIs (191203_CROPt_collapse_by_UMIs.cpp, run within 191203_CROPt_Step2_collapse_UMIs_wrapper.txt) and aligned to the reference sequence using needleall (http://emboss.sourceforge.net/apps/release/6.6/emboss/apps/needleall.html) with default settings. To remove PCR amplification or sequencing errors being interpreted as a CRISPR edit, we devised a strategy to disentangle likely editing from technical errors in sequences where indels or mismatches appeared discontinuous and/or did not overlap the CRISPR cut site. Beginning at the cut site and moving in either direction, each part of a real “edit” had to be within 4 bases of the last position of an edit. This reduces the possibility that a technical error will be counted towards an edit, while allowing for some edits which appear discontinuous. These likely result from complex events in which bases were both deleted and inserted, with small fragments of insertions mapping to the reference sequence of the deleted region.

Editing at each target in each cell was then evaluated. An unambiguous target was defined as one which either contained no discrepant editing patterns, or if multiple editing patterns were observed, had more than one UMI (unique transcript) associated with the “real” editing pattern, and no more than 1 of the other (assumed to be either a stray transcript picked up during processing or a product of template switching during PCR). If more than one edit was associated with more than one UMI, the target was termed “ambiguous.” If each edit was only associated with one UMI, the target also was termed “ambiguous.” For the two duplicated targets, if ambiguous editing patterns were distributed in silico as described below.

The above steps are implemented in wrapper script 191205_local_target_analysis_all_UPDATED.txt.

### Evaluating CRISPR target capture rates and filtering cells based on target capture and expression

The dual sci-RNA-seq + target capture was performed in eight batches. The median number of targets captured varied by batch (**Supplementary Figure 2**). This discrepancy was traced to the batch of Tn5 buffer used in each batch: more recently made batches as well as the commercial batch (as opposed to older buffer made internally) produced more efficient Tn5 integration into cDNA (readily observed in difference of sci-RNA-seq median library size). Since Tn5 integration occurs prior to separating the samples for separate RNA and target processing, a smaller cDNA fragment size means that Tn5 is more likely to integrate within a target region (downstream of the 5’ primer binding site), thus preventing that target from being captured. Thus, optimization of buffer composition might address this issue.

To filter out presumed doublets, both target editing and expression data were used (**Supplementary Figure 2**). Cells were called “Singlet” of “Doublet” based on fraction of ambiguous targets (those with more ambiguous than non-ambiguous targets were considered doublets). For doublet cells, the sci-RNA-seq UMI count distributions were shifted, indicating that high count cells are likely doublets. In addition to removing cells defined as doublets by target editing patterns, we thus additionally removed cells which were above 1.8x the median sci-RNA-seq UMI count for each batch (**Supplementary Figure 2c**).

### Tree-building algorithm steps

#### (1) Computationally split duplicated targets

Two targets (#30 & 31) were consistently associated with two editing patterns within a single cell, strongly suggesting that the section of chromosome on which these targets reside underwent a duplication event in an early cell division (or in an ancestor of the founder cell of this population). Because editing patterns at these targets clearly contained early editing events which were informative of tree structure, we decided to computationally split each target into two separate targets. For each target, we first generated a list of pairs of edits which were commonly found together in a single cell. Since these had to have occurred at two different targets, we constrained a set of editing patterns to one target and a set to the other. Editing patterns which were frequently found alongside an unedited target (indicating that just a single target of the pair was editing in that subset of cells) or on their own (indicating no duplication or a loss of the duplicated target) were randomly assigned to the first target of the pair. Thus, a list of allowed “edits” was generated for each target in the pair. If a cell contained edits on either list, they were distributed accordingly between the pair of targets. The final dataset thus contains a total of 33 targets per cell.

#### (2) Infer missing data

While a subset of missing data reflects true loss of either the target itself (due to a large deletion or a CNA) or an editing pattern which makes the target hard to capture (e.g. a very large insertion), some targets are stochastically not captured during mRNA processing. We thus attempted to infer these edits using a nearest neighbors approach. Since batch 1 had the most complete lineage data, for correcting missing data from other batches we combined them with batch 1 cells and performed the following steps. We first calculated similarity scores between every pair of cell lineage profiles using an additive approach. For each target with matching editing patterns a score of 5 would be added to the total; for each target that was unedited in both lineage profiles, a score of 1 would be added. Targets which did not match (or contained missing or ambiguous data in either cell) received a score of zero. Based on these similarity scores, we defined a set of “nearest neighbor” cells for each cell, and used these to computationally infer missing data for each cell. Specifically, for each cell, for each missing or ambiguous target, we used the most common editing pattern in its closest set of neighbors at that target to infer the missing edit. If the majority of neighbors also had missing data at this target, this likely reflects a true loss at this target, and thus was left uncorrected.

Steps 1 & 2 above are implemented in 200713_wrapper_for_wrapper_for_AMBcorr_Xcorr_step.txt.

#### (3) Generate initial groups of related cells using hierarchical clustering

We generated a similarity matrix using the similarity score described above, and hierarchically clustered cells via Ward’s method (Ward2 in “hclust” package in R). Duplicated targets described in “Computationally split duplicated targets” (targets 30-33) were not used for similarity calculations as they were found to bias groupings. Trees generated via hierarchical clustering are not consistent with progressive CRISPR-based editing events, but do a reasonable job of placing similar cells next to one another. Hierarchically clustered trees can be split automatically into a desired number of groups, but we found that for downstream applications, it was best to manually determine how to split the tree since in some cases groups of very different sizes were desired. We thus generated plots of the hierarchically clustered tree (resembling the inset in **Figure 3e** but containing the full tree) and manually chose the break points at which groups should be split. We generated plots of both the lineage profile in which we had inferred missing data as in step 2, and of the raw data, and consulted both to ensure missing data inference appeared accurate. Importantly, these groups were chosen with the intention that some would be split further in a subsequent step: as long as cells appeared confidently as close relatives, they were kept in a single group at this stage. This procedure generated 94 groups. Groups with less than three cells were removed to be placed into larger groups at a subsequent step, leaving 45 groups remaining.

Groups were evaluated visually as implemented in 200811_combine_like_cells_for_loop.R, 200219_make_LG_group_plots_for_combined_cell_groups.R, & 200225_plot_many_LG_on_one_plot_from_Refcell_list.R.

#### (4) Generate a “consensus” lineage profile for each group

A consensus editing pattern at a target was defined as one which appeared in at least 75% of cells in that group. A single consensus lineage profile was first generated automatically using this definition for each group. We then manually corrected these profiles to account for known sources of missing data which may contribute to an editing pattern being captured at fewer than 75% of cells. For example, large insertions and deletions are captured less efficiently, and thus a target in which contains >25% of missing data, but the remaining cells contain a consistent large insertion or deletion, we can plausibly infer that that editing pattern is likely present in all cells.

#### (5) Generate a preliminary lineage tree of consensus cells via an iteratively applied greedy approach

If no data were missing and no convergence (identical edits occuring at a single target independently) were present, one could theoretically build a perfect tree using the greedy approach shown in **Figure 3c**. First, we identify the most abundant editing pattern at a single target in the tree, and split the consensus cells into two groups based on the presence or absence of this editing pattern. This defines the first branch point. We then apply this approach to the two new subgroups, and iteratively apply it to all subsequent groups to generate a bifurcating tree with leaves being defined by a single consensus lineage profile (implemented in 201109_building_a_tree_3_record_all_changes.cpp). We then collapse any bifurcations which are not supported (when a branch is formed which is not defined by a specific editing event), such that greater than two branches can arise from a single node (201109_AUTO_collapse_bifurcations.R).

Though the consensus editing patterns are not perfect with regards to the above algorithm (there are several instances of convergence, and some missing data), the pooling of related cells to increase confidence of consensus editing patterns makes the algorithm above a viable approach. We thus applied it to the preliminary group of consensus lineage profiles to generate a preliminary tree.

As described above, some groups could be subdivided further. We thus applied the above algorithm to subgroups of the tree, by taking all cells within a single consensus lineage profile, subdividing them into smaller “consensus” groups (beginning with hierarchical reclustering), and generating a subtree as described above. These subtrees were then combined to form the larger tree.

Importantly, this approach of successive tree and subtree generation allows us to deliberately leave out potentially problematic targets, and to choose different sets of targets for each subtree reconstruction. For example, since targets 30-33 contained missing data which may have been the product of edit pattern distribution to resolve target duplication, we removed these for the initial hierarchical clustering which generated cell groups, but used this information for consensus lineage profile calling and greedy tree generation.

Though branching order correctly describes the order of editing events, the depth of the branching events shown in **Figure 3e** does not necessarily indicate a true temporal relationship. Depth on the tree correlates with the number of edits which occurred over the course of that branch’s formation but should not be interpreted as temporal relationships as a consistent editing rate cannot be assumed.

#### (6) Visualizing preliminary trees for manual correction of missing data and resolution of convergence events

Visualizing these trees at various stages allowed us to refine the trees further by helping to resolve previously unclear editing patterns within some consensus cells. For example, the edit at target 26 in groups 33-42 is a large insertion which is not efficiently captured. The majority of cells within groups 33-40 contained missing data at this target, while a subset contained the insertion. But based on the edit in target 31, it appears most likely that all cells actually did contain the insertion at target 26, but it was not captured well. We thus manually corrected targets at which events like these appeared to be the case.

Visualization of intermediate trees also helped to resolve convergence events. Though few convergence events (defined as the same edit occurring multiple times independently at the same target) impacted the automatically-generated tree structure as earlier subdivisions isolated these events from one another, this was not the case in a few places in the tree. In these cases, a group which visually appears to be closely related to another group because of subsequent shared edits is separated from it in early divisions. These events were manually corrected as well.

In two instances, several convergence events were also resolved by shared CNAs between groups. This was rare; with the exception of the instances described below, expression data was not used for tree reconstruction.

Change 1: A single discrepancy (copy number pattern on chromosomes 5 & 11) revealed a convergence event whereby a common editing pattern occurring independently (target #7, teal edit) forced groups together improperly. Instead, a common CNV pattern at chromosomes 5 & 11 strongly suggested that groups 16-19 shared a common ancestor. A change was made accordingly, slightly increasing tree resolution.

Change 2: CNVs on chromosomes 6 & 11 also allowed for better resolution of groups 38-42, where a combination of factors including a convergence event of a commonly observed edit and a large insertion event frequently manifesting as missing data made it challenging to resolve tree structure.

We found for downstream analysis that small groups reduced power below the level at which meaningful expression and accessibility differences could be detected. We thus recombined some closely related groups such that the minimum number of cells per group is 34.

In the end, the final tree contained 42 lineage groups.

#### (7) Integrating remaining cells into pre-defined consensus lineage groups

About a quarter of the cells (batches 1 & 3) were used to construct the original tree. Some of these which formed a group of 1 or two cells in step 3 were removed to be placed into larger groups later, along with the remaining three quarters of the cells w/ lineage profiles. We placed cells into their most closely related groups by calculating similarity scores described above(see (2) <u>Inferring missing data</u> above) on uncorrected lineage profiles with cells already in the tree, and placing new cells into the group in which they had the highest similarity scores. If a cell had identical similarity scores w/ cells from multiple groups, it was placed into the group in which it had the most neighbors.

Final lineage groups were evaluated visually, by plotting lineage profiles of all cells in a single group and visually confirming shared editing patterns.

#### Tree lineage profile visualizations

Tree visualizations were generated using custom code (200807_AUTO_tree_custom_visualization_organized.R, internally running 200806_make_coordinates_for_tree_plot.cpp), which converted tree structure into line segment coordinates which can be plotted in a ggplot space alongside visual lineage profiles. Input files are provided (tree_file_LinRNA, lineage_profiles_wRNA.txt).

Visualizing single lineage groups (**Figure 7c**) implemented in 211129_CopyForFigsRNA_Uncorr_AUTO_tree_custom_visualization_organized.R (RNA) & 211129_CopyForFigsATAC_Uncorr_AUTO_tree_custom_visualization_organized.R (ATAC).

### Permutation Analysis for DE gene identification

#### DE genes were identified using the following procedure

First, raw counts were scaled to 10,000 reads per cell. Then, for each pair of sister groups within the tree (defined as those that share an immediate common ancestor branch), cells were permuted into two groups of the original sizes 10,000 times and the log-fold change for each gene was calculated. Only genes which were expressed in at least 10% of cells in either group were kept for downstream analysis. The measured (real) mean expression ratio for each gene was ranked against the permuted values, for a total of 10,001 values. Z-scores are calculated here as the distance of the real log ratio from the mean divided by the standard deviation of the permuted values.

To account for differences in group sizes across the tree, as well as large CNVs, we evaluated genes on each chromosome in each pair of groups separately to determine the rank cutoff values associated with significant DE. We chose a false discovery rate cutoff of 5%.

Rank cutoff values for each chromosome-group pair combination were determined as follows. If no genes on a chromosome were differentially expressed, we would expect a uniform rank distribution for 1 to 10,001. Thus, the expected number of genes observed at any given rank value is the total number of filtered genes on chromosome/10,001, referred to here as the baseline value. If true DE genes are present, we should observe an enrichment of genes at either or both ends of the distribution, manifesting as higher counts and denser coverage.

An FDR value for each rank position can be determined simply by subtracting the baseline value from the total gene count at each rank. Since those genes of rank 1 or 10001 are most likely to be true positives, we begin at the ends and move inward to identify a group of ranks which together produce an FDR of <= 5%.

The procedure to determine significant ranks is implemented as follows. We begin at rank 1 or 10001, choosing the one with the highest observation count, and calculate the FDR associated with that rank. If it is smaller than 5%, we compare the next most extreme ranks (2 or 10001 if rank 1 was already used), and again choose the one with the highest gene count. We calculate the total FDR encompassing both rank positions and continue this procedure iteratively, until the FDR reaches 5%. All genes with the ranks identified by this procedure are considered DE.

Genes which were lowly expressed in both groups being compared (defined as those for which the percent of cells expressing the gene, calculated separately and then summed between the two groups, is <10%) were removed from the final analysis.

Procedure implemented in A_210327_perm_qsub_script.sh & 210330_process_permutation_table_log_version.cpp.

#### sci-RNA-seq visualization

For heatmap plotting, counts per gene were pooled by lineage group, and a mean was calculated for each gene using the total number of cells as the denominator. Genes with low total counts across the dataset were removed. Specifically, a lowly expressed gene was defined as one which was expressed at a mean of .5 counts per cell or less in all lineage groups. For each retained gene, the lineage group mean was divided by the mean expression in all cells of that gene, and a log2 was taken to center around 0. For visualization scaling purposes, values above or below 1 & -1 (Figure 4) and 1.5 and -1.5 (all other figures), respectively, were changed to those values. Visualization implemented in 201117_AUTO_NewGroups_BETTER_long_AllChr_heatmap_plot.R. Pileups were plotted using ArchR (Granja et al., 2021).

#### SNP-based copy number analysis

To identify variable genomic positions from expression data, a 4 column file was generated for each chromosome from the STAR alignment output file, including cell name, mapping position, CIGAR string, and sequence, and the frequency of each base was calculated as implemented in 201114_wrapper_for_ASEs_for_lineage_groups.txt. Counts were generated for all cells as well as subsets of groups. Variable positions were retained and SNP info was added to via code 191018_add_snp_info_to_ASE_file.cpp, using as input a tab-delimited file generated from a vcf file, containing five files: chromosome, position, rs_id, major allele, minor allele. Plots were generated in 200203_ASE_calc_major_freq.R.

#### sci-ATAC-seq processing

For processing sci-ATAC-seq sequencing reads, we first compare observed and expected lists of single cell indices, correcting any indices with a likely off-by-one error. All reads are then adaptor trimmed using trimmomatic (parameters: TRAILING:3 SLIDINGWINDOW:4:10 MINLEN:20), and all reads associated with a single cell are then aligned to the genome using bowtie2 (hg38 genome build). Reads are then deduplicated by UMIs using a custom script (191226_CROPt_process_atac_bedfile.cpp). Both cell by gene and cell by interval counts were generated using a custom script (191226_CROPt_make_cell_by_interval_count_file.cpp). During analysis, count files were converted into the 10X Genomics format for compatibility with other analysis tools.

For heatmap plotting, counts per gene/interval were pooled by lineage group, and a mean was calculated for each gene using total number of UMIs (as opposed to total number of cells) as the denominator to account for a large spread of total observed UMIs per cell. Each value was then scaled by the median of the total read count for all genes/bins. Genes & bins with low total counts across the dataset were removed (those whose scaled values were below 120 per 1MB bin, or below 5 per gene, in all groups). For each retained gene/interval, the lineage group mean was divided by the mean accessibility of all cells at that gene/interval, and a log was taken to center around 0. For visualization scaling purposes, values above or below .9 & -.9 respectively were changed to those values. This was implemented in 210222_ATAC_process_bin_counts_by_groups_play_w_scaling.R.

Differential accessibility was evaluated using the permutation approach described above, with mean counts per a group again calculated with total number of UMIs (as opposed to total number of cells) as the denominator.

Pileups were plotted using ArchR (Granja et al., 2021). For DA analysis at peaks, a set of peaks was determined using ArchR, using both the whole dataset as well as successive subgroups moving across the tree. The union of these peaks was then overlapped with DE genes (including 5kb upstream) and DA at these peaks was again evaluated using the permutation approach.

## Data & Code Availability

Raw and processed data and code are available on GEO (GSE201339) & Github (https://github.com/minkinaa/TheLorax). See README on Github for further details.

## Acknowledgements

We thank Darren Cusanovich, Riza Daza, Jean-Benôit Lalanne, Jonathan Packer, and Erick Matsen for helpful feedback on experimental protocols and analysis approaches. We thank the members of the Shendure Lab, as well as members of the Allen Discovery Center for Cell Lineage Tracing, for helpful discussions. This work was supported by the NSF Graduate Research Fellowship Program (A.M., 2017-2021), an NHGRI Genome Training Grant (T32) (A.M.), the Paul G. Allen Frontiers Group (Allen Discovery Center for Cell Lineage Tracing to J.S.), the National Institutes of Health (UM1HG011586 and UM1HG011586 to J.S.). J.S. is an Investigator of the Howard Hughes Medical Institute.

## Supplementary Figures

**Supplementary Figure 1.**
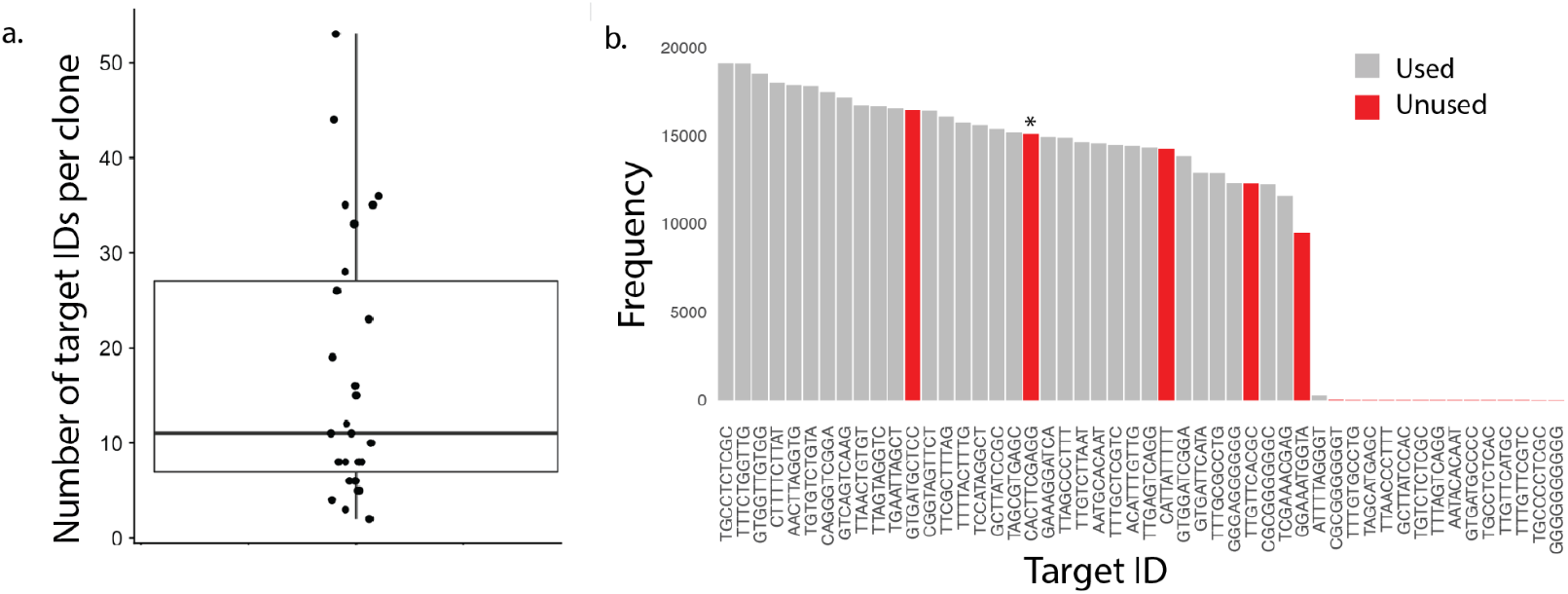
Evaluating lentiviral target integrations. (**a**) Number of unique target IDs across 26 clones derived from high MOI transduction of HEK293 cells. Box shows median and encompasses counts in the second and third quartiles. Whiskers depict the interquartile range. (**b**) Frequency of each unique target ID within the unedited clone used for the main experiment. As discussed in the text, this clone was “re-cloned” following transduction with doxycycline-inducible Cas9 lentiviral construct, such that a single founder cell generated the tree. Four target IDs that were abundant after the first round of cloning were unobserved after this re-cloning step (red bars), while an additional one was corrupted by a mutation and therefore also excluded (red bar with asterisk). The remaining 31 abundant target IDs were carried forward in the analyses, with two of these “duplicated” *in silico* to account for their inferred duplication just before or during the clonal expansion.

**Supplementary Figure 2.**
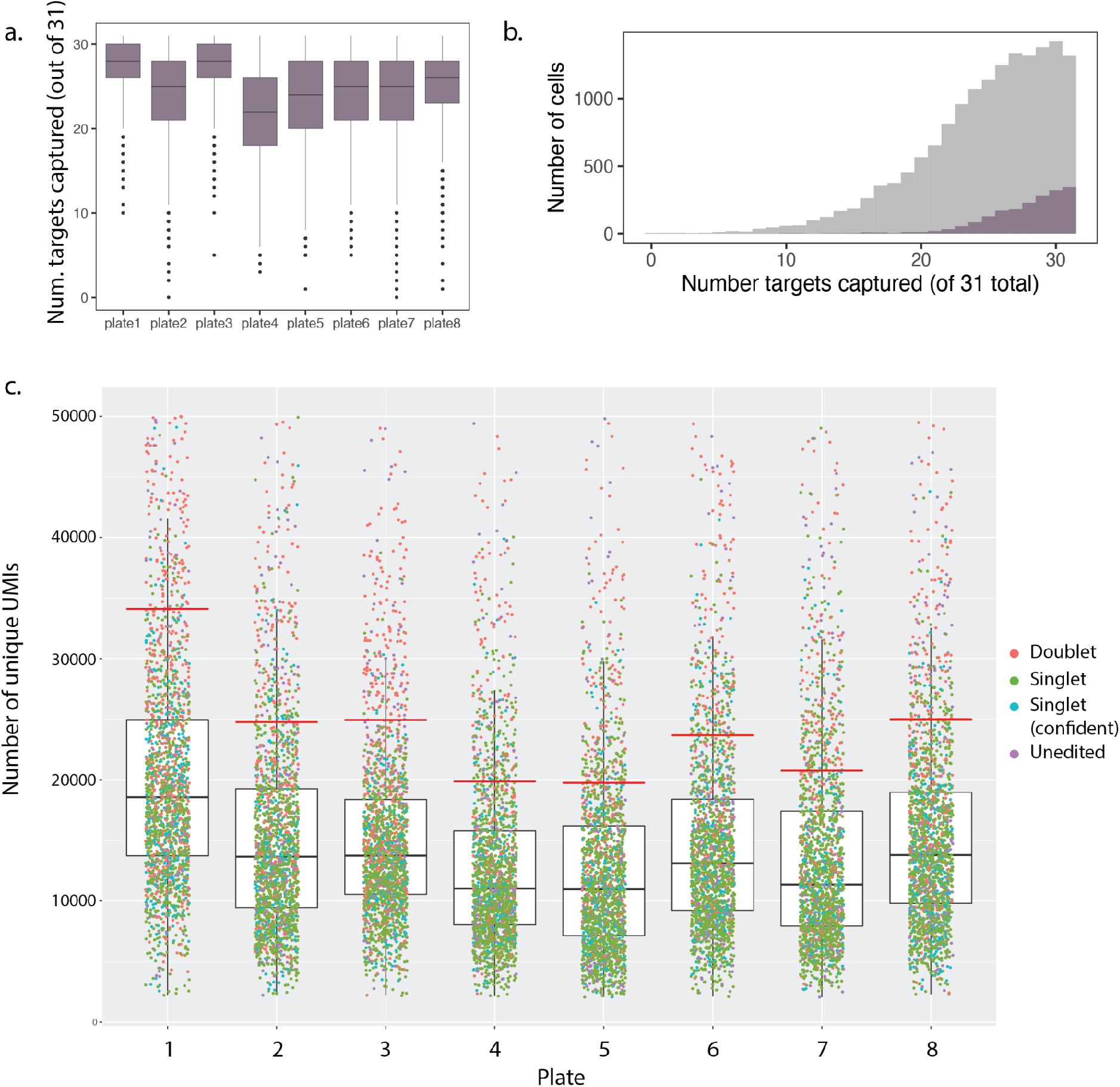
Batch-specific evaluation of target capture. (**a**) Distribution of the number of targets captured per cell, per batch (out of 31). (**b**) Gray: Number of targets captured per cell across batches; Purple: number of targets captured per cell in batch #1. (**c**) Distribution of transcriptome UMIs per cell, per indexed PCR batch (“plate”), with UMI cut-off for doublet removal shown by red lines. Cells with UMI counts > 1.8X the median UMI count for each batch were removed from the analysis. Singlets and doublets inferred from collisions in lineage data. “Singlet (confident)” corresponds to cells which can confidently be called as singlets based on the number of non-ambiguous editing events observed.In panels **a & c**, boxes show median and encompass counts in the second and third quartiles, while whiskers depict the interquartile range.

**Supplementary Figure 3.**
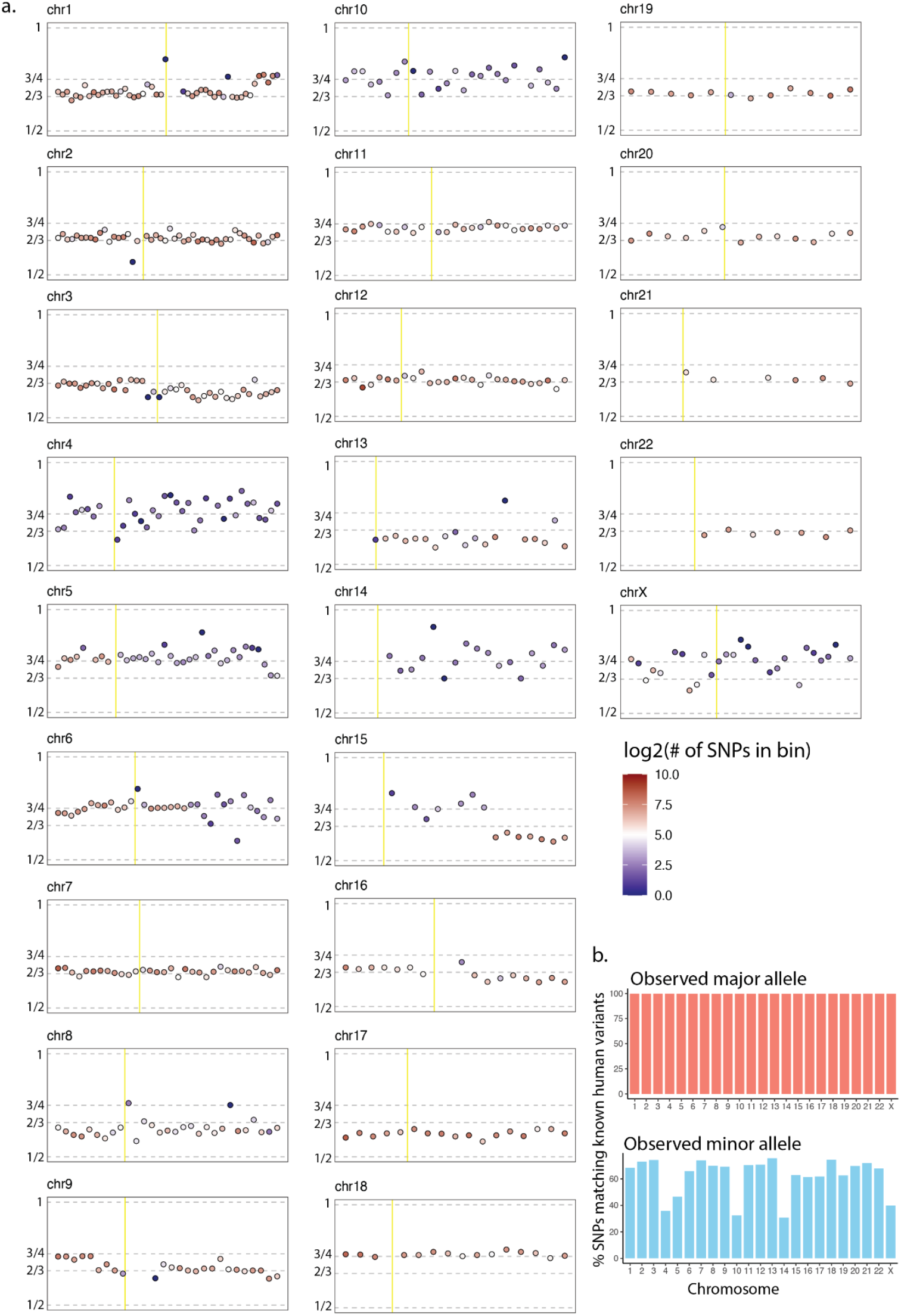
Allelic-ratio-based copy number analysis for all chromosomes. (**a**) Analysis described in Figure 5a-b, performed on all cells for all chromosomes. Point fill color represents the number of SNPs found to be heterozygous in that bin, signaling the reliability of this analysis at that location. Yellow line indicates the centromere position. (**b**) Percent of inferred major and minor alleles at variable positions in the data (filtered as described in **Figure 5a**) which match SNP bases found in humans at those positions (dbSNPs). For simplicity, only single-base SNPs with at most two common alleles in the population were considered.

**Supplementary Figure 4.**
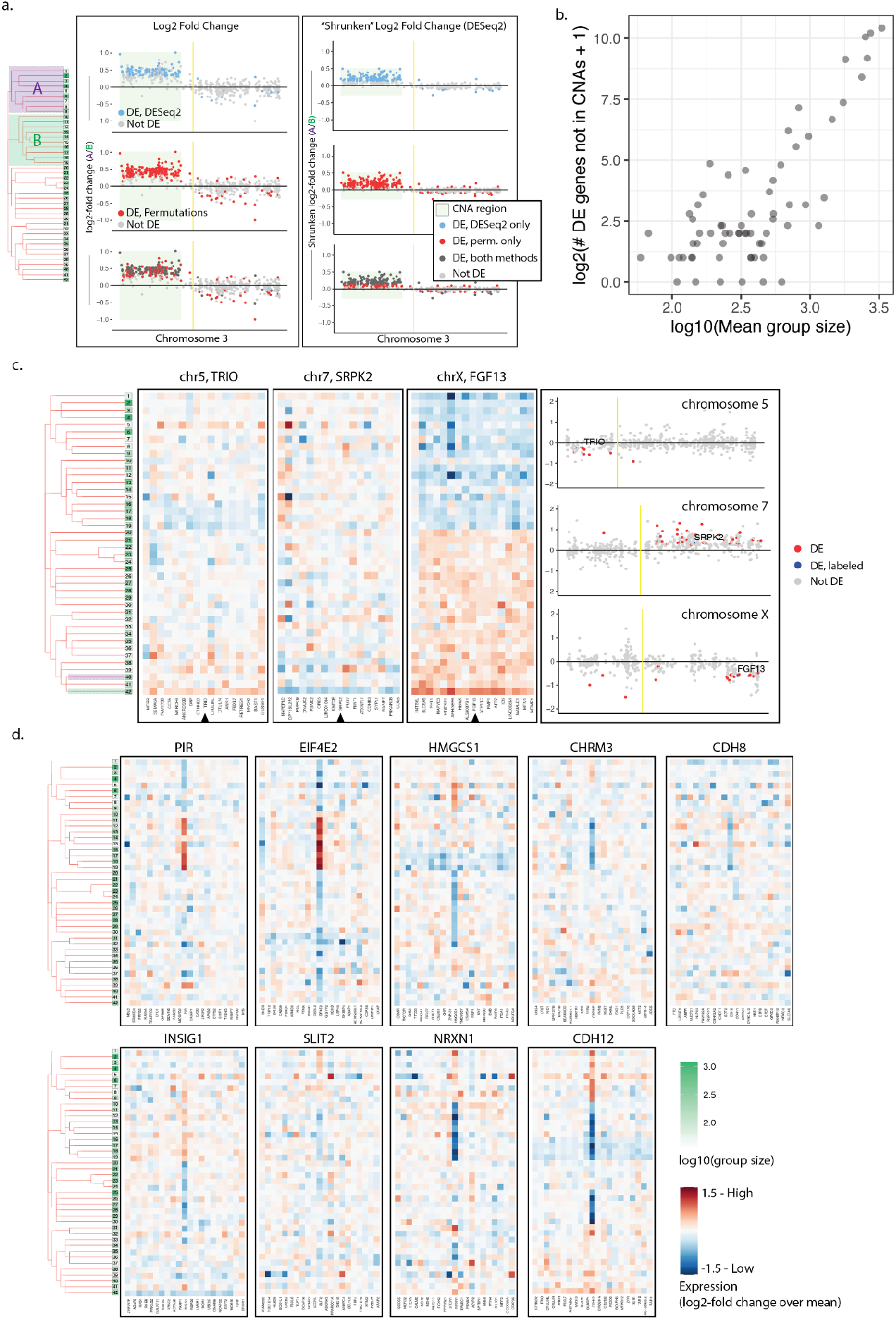
Differentially expressed genes within and outside of detected CNAs observed across sister lineage group comparisons. (**a**) DE genes detected by the permutation approach vs. DESeq2. The left plots show log2-fold changes, while the right plots show the “shrunken” log2-fold changes calculated by DESeq2, which takes absolute expression level into account, and corrects for higher variance at low expression levels. (**b**) Relationship between group size (mean of the two groups being compared) and DE genes not associated with a CNA. (**c**) DE genes detected within CNA regions on chrs 5, 7, and X, between the indicated groups (234 and 276 cells, respectively). (**d**) Heatmaps showing single genes (middle of each plot) which exhibit heritable expression patterns consistent with the tree structure. Surrounding genes are not DE, suggesting these patterns are not due to CNAs, although we cannot rule out highly focal amplifications with gene expression data alone.

**Supplementary Figure 5.**
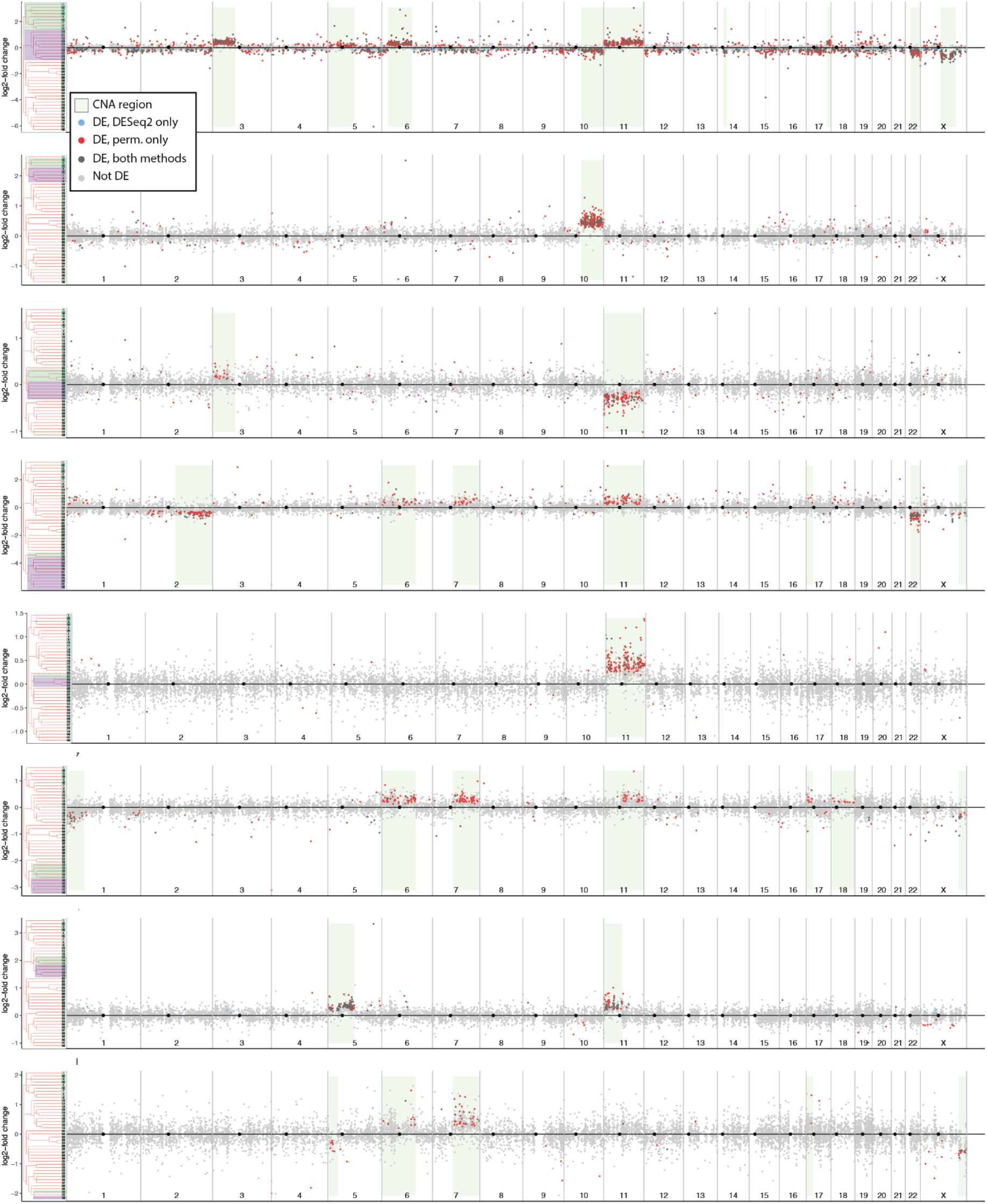
Global DE between select pairs of sister groups. Log2-fold change for expressed genes across all chromosomes between select pairs of sister lineage groups. Groups that are compared in each plot are indicated on the trees at the left with green and purple boxes. Colors indicate by which method (if any) a gene was found to be differentially expressed. Inferred CNAs are shown as light green boxes.

**Supplementary Figure 6.**
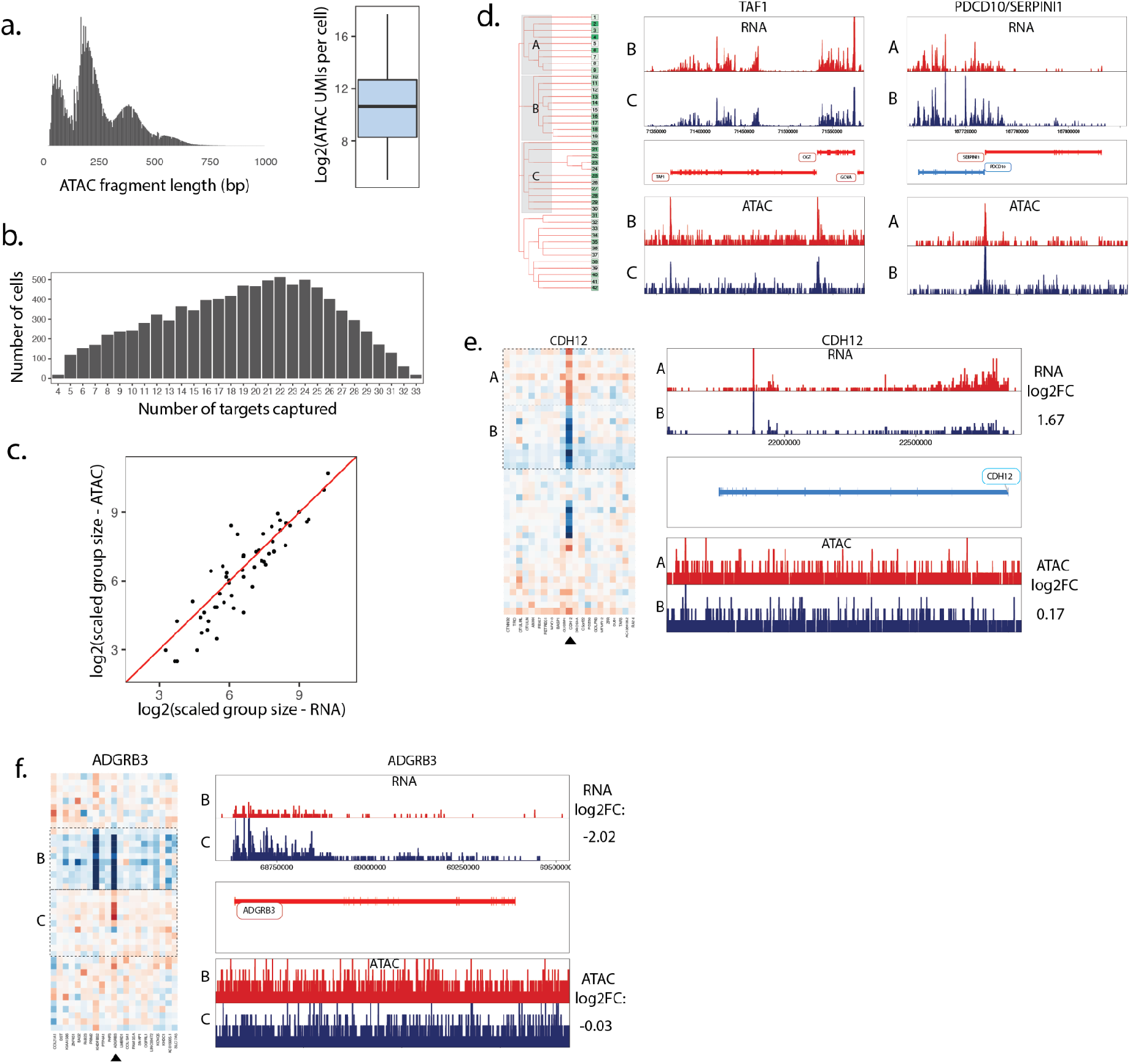
Evaluating lineage-linked chromatin accessibility and expression. (**a**) Histogram of sci-ATAC-seq fragment lengths across all cells (left) and a boxplot of sci-ATAC-seq reads per cell (right). (**b**) Histogram of the number of targets captured per cell included in the analysis. (**c**) Correlation of group sizes collected along sci-RNA-seq and sci-ATAC-seq. Each point represents a single lineage group. Group sizes were normalized to a total cell count of 10,000 for each feature. (**d**) Read pileups for RNA (top) and ATAC (bottom) data for the lineage groups and genes indicated on the tree. Associated heat maps shown in **Figure 7e**. (**e**) Left: Heatmap of relative expression of *CDH12* and surrounding genes Right: Pileup of expression and chromatin accessibility data for the indicated groups (as labeled on tree in **Figure 7g**) at the *CDH12* locus. (**f**) Same as panel **e**, but for *ADGRB3*.

